# Regulated release of cryptococcal polysaccharide drives virulence and suppresses immune cell infiltration into the central nervous system

**DOI:** 10.1101/186668

**Authors:** Steven T. Denham, Surbhi Verma, Raymond C. Reynolds, Colleen L. Worne, Joshua M. Daugherty, Thomas E. Lane, Jessica C. S. Brown

## Abstract

*Cryptococcus neoformans* is a common environmental yeast and opportunistic pathogen responsible for 15% of AIDS-related deaths worldwide. Mortality primarily results from meningoencephalitis, which occurs when fungal cells disseminate from the initial pulmonary infection site and spread to the brain. A key *C. neoformans* virulence trait is the polysaccharide capsule. Capsule shields C. neoformans from immune-mediated recognition and destruction. The main capsule component, glucuronoxylomannan (GXM), is found both attached to the cell surface and free in the extracellular space (as exo-GXM). Exo-GXM accumulates in patient serum and cerebrospinal fluid at μg/mL concentrations, has well-documented immunosuppressive properties, and correlates with poor patient outcomes. However, it is poorly understood whether exo-GXM release is regulated or the result of shedding during normal capsule turnover. We demonstrate that exo-GXM release is regulated by environmental cues and inversely correlates with surface capsule levels. We identified genes specifically involved in exo-GXM release that do not alter surface capsule thickness. The first mutant, *liv7∆*, released less GXM than wild-type cells when capsule is not induced. The second mutant, *cnag_00658∆*, released more exo-GXM under capsule-inducing conditions. Exo-GXM release observed *in vitro* correlated with polystyrene adherence, virulence, and fungal burden during murine infection. Additionally, we find that exo-GXM reduces cell size and capsule thickness in capsule-inducing conditions, potentially influencing dissemination. Finally, we demonstrated that exo-GXM prevents immune cell infiltration into the brain during disseminated infection and highly inflammatory intracranial infection. Our data suggest that exo-GXM performs a different role from capsule GXM during infection, altering cell size and suppressing inflammation.

**Importance:** *Cryptococcus neoformans* is a leading cause of life-threatening meningoencephalitis in humans. *C. neoformans* cells produce an immunosuppressive polysaccharide, glucuronoxylomannan (GXM), that is the main component of a protective surface capsule. GXM is also released free into extracellular space as exo-GXM, although the distinction between cell-attached GXM and exo-GXM has been unclear. Exo-GXM influences the outcome of infection, is the basis for current diagnostic tools, and has potential therapeutic applications. This study increases our basic understanding of the fungal biology that regulates polysaccharide release, suggesting that the release of cell-attached GXM and exo-GXM are distinctly regulated. We also introduce a new concept that exo-GXM may alter cell body and capsule size, thereby influencing dissemination in the host. Finally, we provide experimental evidence to confirm clinical observations that exo-GXM influences inflammation during brain infection.

## Introduction

*Cryptococcus neoformans* is a globally distributed saprophytic fungus found associated with certain species of trees and bird droppings (1). However, in immunocompromised humans *C. neoformans* acts as an opportunistic pathogen. Cryptococcal infections are responsible for 15% of acquired immune deficiency syndrome (AIDS) related deaths worldwide, with most cases occurring in sub-Saharan Africa and Asia (2). Due to its global environmental distribution, human exposure to *C. neoformans* is almost universal (1, 3). Infections begin when inhaled fungal spores or desiccated yeast cells enter the lungs, where they are either cleared by the immune system, or contained and persist for a decade or more (4). Upon patient immunosuppression, *C. neoformans* cells can disseminate from the lungs to basically any organ in the body (5). *C. neoformans* proliferates particularly well in the brain, resulting in life-threatening meningoencephalitis (6). In fact, cryptococcal meningoencephalitis is a primary cause of death among HIV-AIDS patients, with mortality rates exceeding 50% in resource poor areas (2).

In contrast to many forms of bacterial and viral meningitis, cryptococcal meningoencephalitis is associated with strikingly low levels of inflammation and infiltrating immune cells into the central nervous system (CNS) of both human patients and mouse models (7-11). This paucity of inflammation is linked to poorer clinical outcomes, and subdued clinical signs that can delay treatment (9, 12, 13).

An essential factor for *C. neoformans* virulence is the conditional production of a thick polysaccharide surface capsule, which can more than double the diameter of a *C. neoformans* cell (14). The primary capsule constituent is glucuronoxylomannan (GXM), which comprises approximately 90% of the capsule mass (15, 16). Surface capsule plays a number of different roles during pathogenesis, protecting *C. neoformans* cells from phagocytosis, complement, and oxidative stress (15, 17, 18). GXM also has numerous immunomodulatory properties that facilitate fungal survival in the host (19). Notably, GXM increases anti-inflammatory cytokine (IL-10) release while dampening proinflammatory cytokine release (IL-12, IFN-γ TNF-α, IL-1B and IL-6) (20-23). GXM disrupts antigen presentation by macrophages and dendritic cells, and can even induce macrophage apoptosis, thereby diminishing T cell proliferation (21, 24-26). GXM can also suppress leukocyte infiltration into sites of inflammation (27-29).

GXM is non-covalently attached to the cell surface during cell surface capsule formation and maintenance (16). It is also found free within the extracellular milieu. This exo-cellular GXM (exo-GXM) reaches mg/mL concentrations in laboratory growth medium (30), and can be observed in the high μg/mL range in patient serum and cerebrospinal fluid (10, 31). GXM serum titers in HIV-associated cryptococcosis patients positively correlate with non-protective immune signatures and increased mortality (32).

Despite longstanding knowledge of the existence of exo-GXM, its connection to cell-associated GXM and the mechanisms behind its release remain largely unclear. One hypothesis has been that exo-GXM is shed mechanically from the cell surface capsule (16, 33). Alternatively, it has been speculated that distinct mechanisms might regulate the production of cell-associated and exo-GXM in response to environmental cues (15, 16, 34). This latter hypothesis is supported by observations that cell-associated and exo-GXM display different biophysical properties (34). Decreased electromobility of exo-GXM under capsule inducing conditions indicates that these differences could occur at the level of polymer length or branching (35-37).

Here we test the hypothesis that exo-GXM production is regulated by environmental conditions. We find that exo-GXM production is inversely related to the thickness of the cell surface-retained capsule and identify genes involved in these processes. Exo-GXM production also correlates with virulence and reduces infiltration of immune cells into the CNS during infection. Together, these data support the idea that exo-GXM plays a critical but distinct role from cell surface GXM during infection.

## Results

### Environmental signals alter exo-GXM levels

To investigate whether exo-GXM release is passive shedding of surface capsule or regulated at some level, we cultured wild-type *C. neoformans* cells for 24 hours under a variety of media conditions. We then measured capsule size and exo-GXM released into the medium. We chose both non-capsule inducing media and a series of capsule inducing media intended to produce a range of capsule induction. We harvested cells, then stained with india ink to measure capsule thickness as the distance from the cell wall to the outer capsule edge **(Fig. 1A)**. We filtered supernatant through a through a 0.22 μm filter to remove cells, then visualized with immunoblotting with the monoclonal antibody (mAb) F12D2 to quantify exo-GXM release as relative staining intensity **(Fig. 1B)**. Exo-GXM band intensities were normalized to yeast nitrogen base (YNB) + 2% glucose levels, which was the condition with the greatest observed levels of exo-GXM.

**Figure 1:**
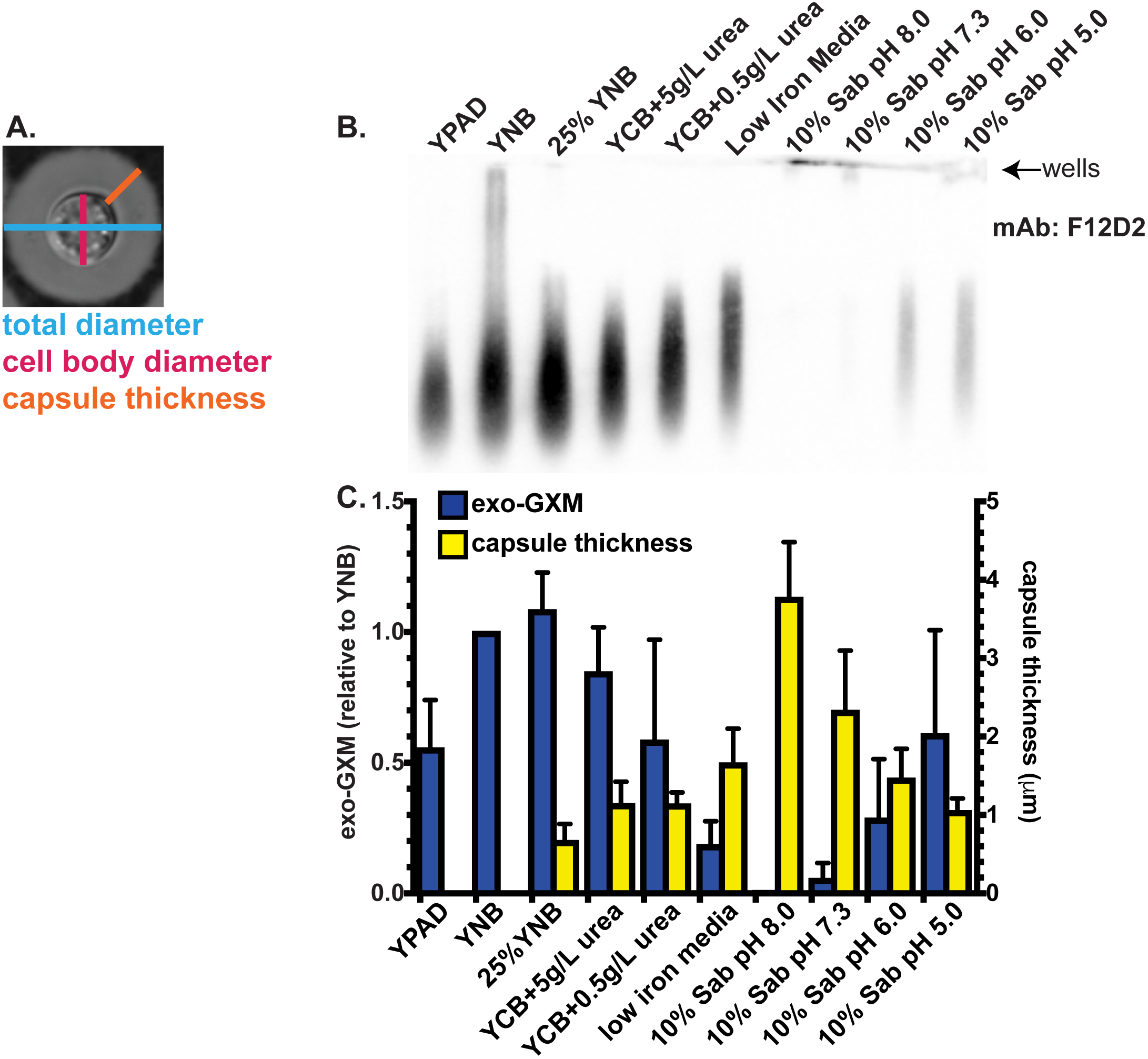
Levels of detectable exo-GXM negatively correlate with capsule thickness under a variety of media conditions. To generate conditioned media, we normalized 24 hour cultures by volume, then passed the supernatants through a 0.22 μm filter to remove fungal cells. **(A)** Representative image of cell/capsule measurements used in this study. **(B)** We tested supernatants for free GXM (“exo-GXM”) by blotting and probing with the F12D2 anti-GXM mAb. See Materials and Methods for further details. A representative blot showing relative levels of exo-GXM collected from cells cultured for 24 hours under a variety of capsule and non-capsule inducing conditions. **(C)** Intensity of exo-GXM bands relative to YNB+2% glucose exo-GXM (blue bars) were quantitated for three independent experiments and plotted next to absolute measurements of capsule thickness (yellow bars) (n=30 cells). Data was combined from three independent experiments. Bars represent the mean and error bars represent the standard deviation (SD).

We found an inverse relationship between capsule thickness and exo-GXM, such that cells growing in the strongest capsule inducing conditions, like 10% Sabouraud’s buffered to alkaline pH, produced the least amount of exo-GXM **(Fig. 1A,C)**. This relationship held across other capsule inducing conditions, such as nitrogen and iron limitation, that produce intermediate levels of both cell surface and exo-GXM.

GXM is an α-1,3-mannan backbone with branching glucuronic acid and xylose residues and variable 6-*O*-acetylation on the backbone (38). *O*-acetylation varies across strains, is not required for capsule formation, but significantly affects GXM’s immunoreactive properties (38-40). Deletion of *CAS1*, which is required for *O*-acetylation, results in a hypervirulent phenotype (41). We analyzed the same conditioned media as in **Figure 1**, but used the mAb 1326 to detect GXM. MAb 1326 recognizes *O*-acetyl (+) GXM, but is unable to recognize *O*-acetyl (-) GXM. F12D2, on the other hand, recognizes both *O*-acetyl (+) and (-) GXM. Thus, 1326 staining intensity relative to F12D2 intensity reflects the relative proportion of *O*-acetyl (+) GXM present in the supernatant. We observed that 1326 staining relative to F12D2 staining increased under certain capsule inducing conditions (low nitrogen, low iron, and 10% Sabouraud’s, pH 5-6), indicative of increased *O*-acetyl (+) GXM **(Fig. S1)**. These results demonstrate that environmental conditions may also influence GXM modification, specifically *O*-acetylation, with potential implications for immune recognition.

### Identification of gene deletion mutants with reduced exo-GXM secretion under non-capsule inducing conditions

We then identified mutants with reduced GXM production. We screened the *C. neoformans* partial knockout collection (CM18 background, 1200 targeted gene knockouts) (42) under YNB, which results in high exo-GXM production. We grew each strain for 24 hours at 37°C, removed the cells by centrifugation, then probed the conditioned medium for exo-GXM.

We searched the YNB-grown mutants for samples that produced less exo-GXM than wild-type cells. We then stained induced cell surface capsule (by growth in 10% Sabouraud’s, pH 7.3) in this subset of mutants and eliminated any with a growth defect and/or a substantial reduction (>25%) reduction in cell surface capsule thickness. We also stained for common pathogen-associated molecular patterns (PAMPs), such as exposed mannoproteins and chitin, which activate host immune responses (43). This left us with a single mutant, *cnag_06464*Δ, or *liv7*Δ, which we re-constructed in the KN99 genetic background (**Fig. 2**). Four other mutants (**Table S1**) exhibited a moderate defect in cell surface capsule in addition to their moderate defects in exo-GXM release. However, we focused on the *liv7*Δ mutant because of its ability to form wild-type levels of cell surface capsule.

**Figure 2:**
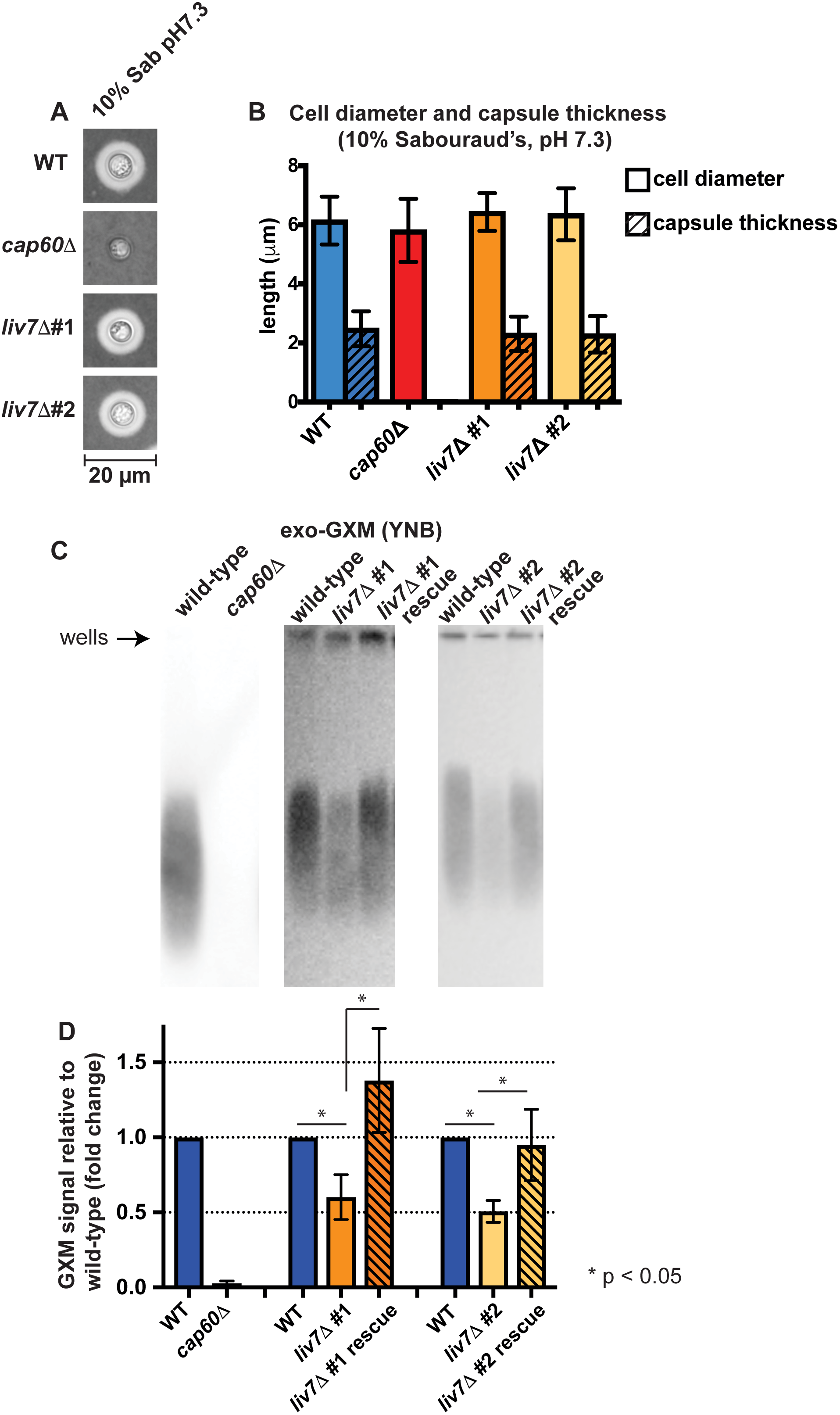
Identification of a genetic mutant (*liv7*∆) with reduced exo-GXM release, but no observable changes to capsule thickness. **(A)** Representative india ink images of cells grown in 10% Sabouraud’s dextrose pH 7.3 for 24 hours. Capsule thickness was similar across KN99 wild-type (WT) cells, and cells from each independent *liv7* deletion strain (*liv7*∆#1 and *liv7*∆#2) **(B)** Quantification of cell body diameter and capsule thickness from three independent experiments (n=30 cells per strain; bars represent mean with SD). **(C)** Conditioned media from wild-type and mutant cultures grown in weak-capsule inducing conditions (YNB + 2% glucose) for 24 hours. Blots were probed with anti-GXM antibody F12D2. **(D)** Quantification of blot signal intensities shows reduced exo-GXM release by *liv7*∆#1 / *liv7*∆#2. Data was combined from three independent experiments. P-values were calculated using a Mann-Whitney test; bars represent mean with SD.

The *LIV7* gene was previously identified in a screen for mutants deficient in growth in the lung (42). Liv7 is localized to the Golgi under capsule-inducing conditions (DMEM + 5% CO_2_) (44). *liv7*Δ cells produce wild-type-like levels of cell surface capsule when grown in 10% Sabouraud’s, pH 7.3 (**Fig. 2A,B**), but conditioned medium from *liv7*D cell cultures grown in YNB contains two-fold less GXM than conditioned medium from wild-type *C. neoformans* cell cultures (**Fig. 2C,D**). PAMP exposure is comparable to wild-type cells (**Fig. S2**).

### Identification of gene deletion mutants with elevated exo-GXM secretion under strong capsule inducing conditions

We next identified mutants that produced elevated levels of exo-GXM under capsule-inducing conditions, when exo-GXM production is very low. We again screened the *C. neoformans* knockout mutant collection (CM18 background), this time growing the mutants in YNB, then subculturing by diluting 1:100 into 10% Sabouraud’s, pH 7.3, and growing 48 hours at 37°C. We again removed mutants that exhibited growth defects, elevated PAMP exposure, and a substantial reduction (>25%) reduction in cell surface capsule thickness. We found two groups of mutants: group #1 exhibited approximately wild-type capsule thickness, while group #2 mutants had less-than-wild-type levels of cell surface capsule (**Table S1**). We focused our subsequent experiments on the mutant in gene *cnag_00658*, which produces cell surface capsule with the same thickness as wild-type cells (**Fig. 3A,B**). As with *liv7*Δ, we re-constructed this mutant in the KN99 genetic background and used those strains for all subsequent experiments. As in the CM18 background, *cnag_00658*Δ cells in the KN99 background released increased exo-GXM in 10% Sabouraud’s, pH 7.3 **(Fig 3C,D)**. Unlike other mutants in group #1, *cnag_00658*Δ cells produce the same levels of melanin and urease as wild-type cells (**Fig. S2**).

**Figure 3:**
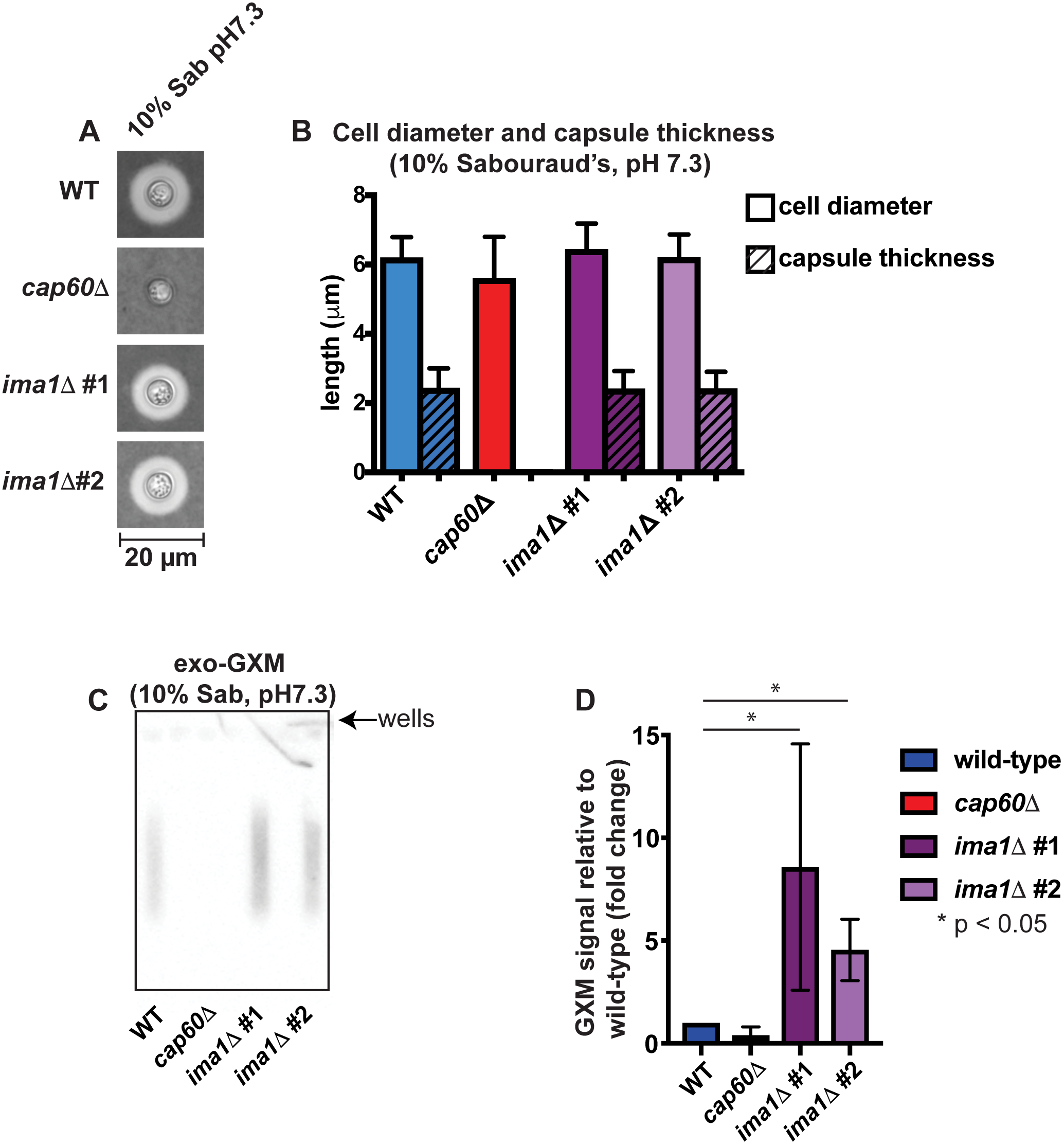
Identification of a genetic mutant (*ima1*∆) with increased exo-GXM release, but no observable changes to capsule thickness. **(A)** Representative india ink images of cells grown in 10% Sabouraud’s dextrose pH 7.3 for 24 hours. Capsule thickness was similar across KN99 wild-type (WT) cells, and cells from each independent *ima1* (also *cnag_00658,* see main text for details) deletion strain (*ima1*∆#1 and *ima1*∆#2). **(B)** Quantification of cell body diameter and capsule thickness from three independent experiments (n=30 cells per strain; bars represent mean with SD) **(C)** Conditioned media from cultures grown for 24 hours under strong capsule-inducing conditions (10% Sabouraud’s at pH 7.3). Blots were probed with anti-GXM antibody F12D2. **(D)** Quantification of blot signal intensities shows increased exo-GXM release by *ima1*∆#1 / *ima1*∆#2 (Combined data from three independent experiments. P-values were calculated using a Mann-Whitney test; bars represent mean with SD.

The *CNAG_00658* gene encodes a predicted protein 624aa in length. It shares N-terminal sequence homology with the *Schizosaccharomyces pombe* inner nuclear membrane protein, IMA1 (615aa). *CNAG_00658*’s predicted gene product also has five putative transmembrane domains that positionally align with the 5 transmembrane domains of the *S. pombe* IMA1 protein. For these reasons, we propose to rename the *CNAG_00658* gene, “*IMA1*”. For the duration of this text, we will refer to “*cnag_00658*” as “*ima1*”.

### Changes in exo-GXM levels alter fungal cell adherence

We had thus far only assayed exo-GXM secretion during planktonic growth. However, within its natural environment of soil and vegetable matter, *C. neoformans* can form adherent biofilms (45). Previous work on cryptococcal biofilms has revealed that a significant portion of the extracellular matrix is composed of GXM, and that it plays a critical role in adherence (46). Acapsular strains are unable to adhere to surfaces such as polystyrene, and the addition of anti-GXM antibodies to developing wild-type biofilms reduces their adherence (46). We speculated that exo-GXM may be incorporated into the extracellular matrix during sessile growth to provide community level structure, and that our exo-GXM mutants would display varying adherence corresponding to their exo-GXM secretion profiles.

To test this, we grew cells at a concentration of 10^6^ cells/100ul in 96 well polystyrene plates at 37°C. After 48 hours, the wells were washed forcefully with PBS+0.1% tween-20 dispensed from an automated plate washer, resuspended in PBS containing XTT/menadione and left for 5 hours at 37°C. XTT is reduced by fungal cells to produce a colorimetric measure of metabolism that is highly correlative with viable cell count (47).

Wild-type, *cap60*∆, *liv7*∆#1 / #2, and *ima1*∆#1 / #2 cells were assayed in both YNB and 10% Sabouraud’s pH 7.3 to replicate planktonic non-capsule and capsule-inducing conditions respectively. The *cap60*∆ cells served as a negative control, as acapsular mutants are unable to adhere, likely due to their lack of surface and exo-GXM (46). We hypothesized that *liv7*∆#1 / #2 cells would display reduced adherence in our assay due to the reduction in exo-GXM release we observed during planktonic growth. This was indeed the case, as we observed an approximately two-fold reduction in the ability of *liv7*∆#1 / #2 cells to adhere in our assay **(Fig. 4A)**.

**Figure 4:**
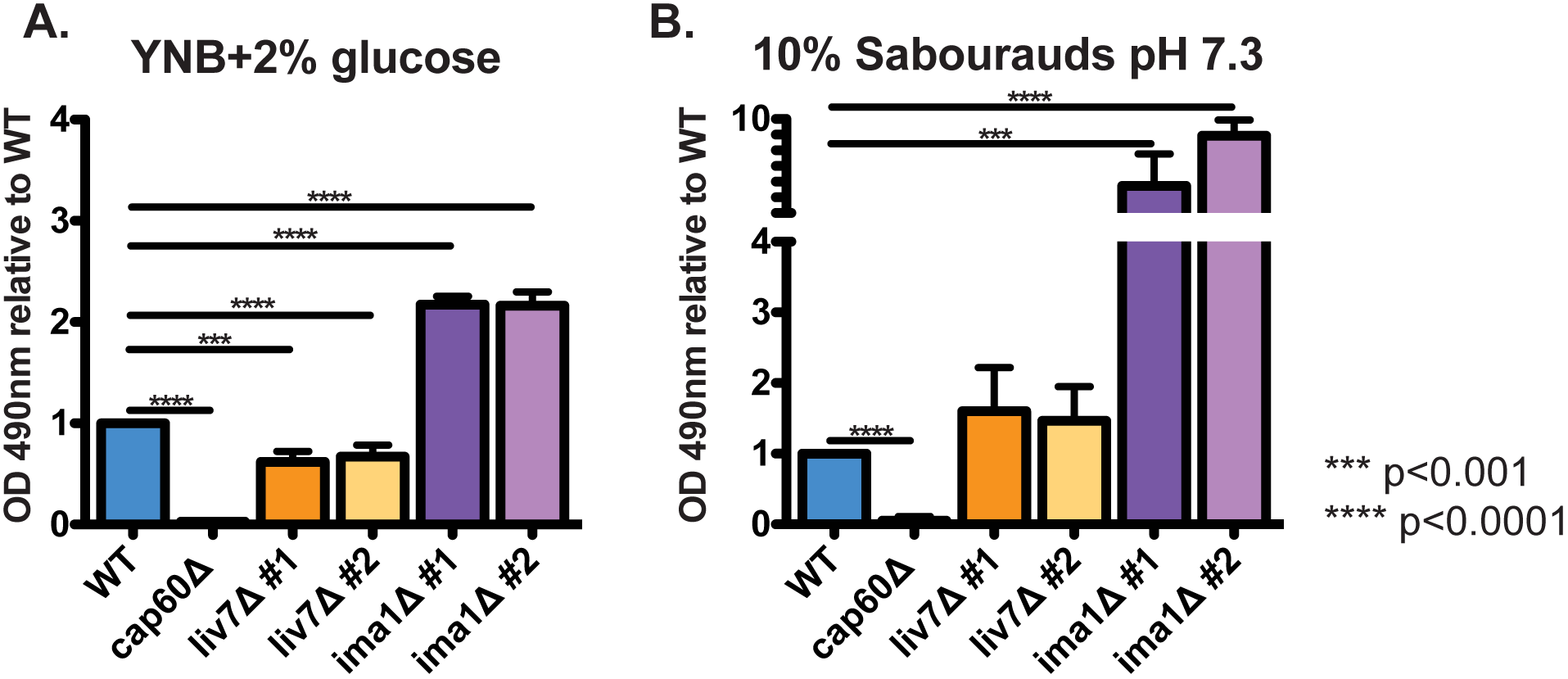
Mutants’ alterations to exo-GXM release correlates with adherence. 10^6^ *C. neoformans* cells were seeded into individual wells of 96-well polystyrene plates and incubated at 37°C. 48 hours later, the wells were washed to remove non-adhered and/or weakly adhered cells before resuspension in XTT for colorimetric analysis of metabolic activity as a proxy for viable cell count. **(A)** OD_490_ readings from cells grown in YNB, normalizing to wild-type cell readings. *liv7*∆#1 / #2 cell adherence was reduced and *ima1*∆#1 / #2 cell adherence was increased when compared to wild-type cells. **(B)** OD_490_ readings from cells grown in 10% Sabouraud’s pH7.3, normalizing to wild-type cell readings. *ima1*∆#1 / #2 cell adherence was increased when compared to wild-type cells. Combined data from three independent experiments. P-values were calculated using a Mann-Whitney test; bars represent mean with SD.

In contrast to YNB, *liv7*∆#1 / #2 cells were able to adhere at wild-type levels when grown in 10% Sabouraud’s pH 7.3, perhaps because our observations of planktonic cells indicated that far less exo-GXM is released by both wild-type and *liv7*∆#1 / #2 cells under these conditions **(Fig 4B)**. Similarly, *ima1*∆#1 / #2 cells, which displayed elevated exo-GXM secretion under strong capsule inducing conditions, demonstrated six to eight-fold higher adherence than wild-type when grown in 10% Sabouraud’s pH 7.3 **(Fig 4B)**. When grown in YNB, *ima1*∆#1 / #2 cells still displayed increased adherence, but it was reduced to an approximately two-fold increase over wild-type **(Fig 4A)**. Altogether, these results suggest that the regulated secretion of exo-GXM may have a specialized role in an environmental setting by promoting the adherence of *C. neoformans* communities.

### Host survival and fungal burden correlates with in vitro exo-GXM levels

Next, we sought to use *liv7*∆#1 / #2 and *ima1*∆#1 / #2 as an opportunity to explore roles for exo-GXM during pathogenesis. We hypothesized that exo-GXM secretion would promote virulence through its immunomodulatory properties. Since *liv7*∆#1 / #2 and *ima1*∆#1 / #2 cells produce wild-type sized surface capsules in culture, we anticipated that *liv7*∆#1 / #2 and *ima1*∆#1 / #2 cells would allow us to assess the role of exo-GXM in pathogenesis, independent of surface capsule. We predicted that the reduction of *liv7*∆#1 / #2 cells’ ability to produce exo-GXM *in vitro* would result in reduced virulence. Similarly, we predicted that *ima1*∆#1 / #2 cells, which show increased exo-GXM under capsule inducing conditions, would display heightened virulence.

We employed a murine model of disseminated cryptococcosis by inoculating C57BL/6NJ mice (Jackson Labs) intranasally with 2.5x10^4^ fungal cells per mouse. We calculated survival as the time it took each mouse to reach 85% of their initial mass. Consistent with our hypothesis, *in vitro* exo-GXM production inversely correlated with host-survival. Wild-type KN99 infected mice reached endpoint a median of 20 days post-inoculation (dpi). In contrast, *liv7*∆#1 / #2-infected mice reached endpoint a median of 22.5 dpi, and *ima1*∆#1 / #2-infected mice a median of 18 dpi **(Fig. 5A)**. However, it is important to note that all strains were sufficiently virulent to cause lethal infection at our inoculating dose. This was not altogether unexpected, as the exo-GXM secretion phenotypes for *liv7*∆#1 / #2 and *ima1*∆#1 / #2 cells were dependent on growth conditions, and manifested as a gradient of exo-GXM production rather than complete ablation or overexpression.

**Figure 5:**
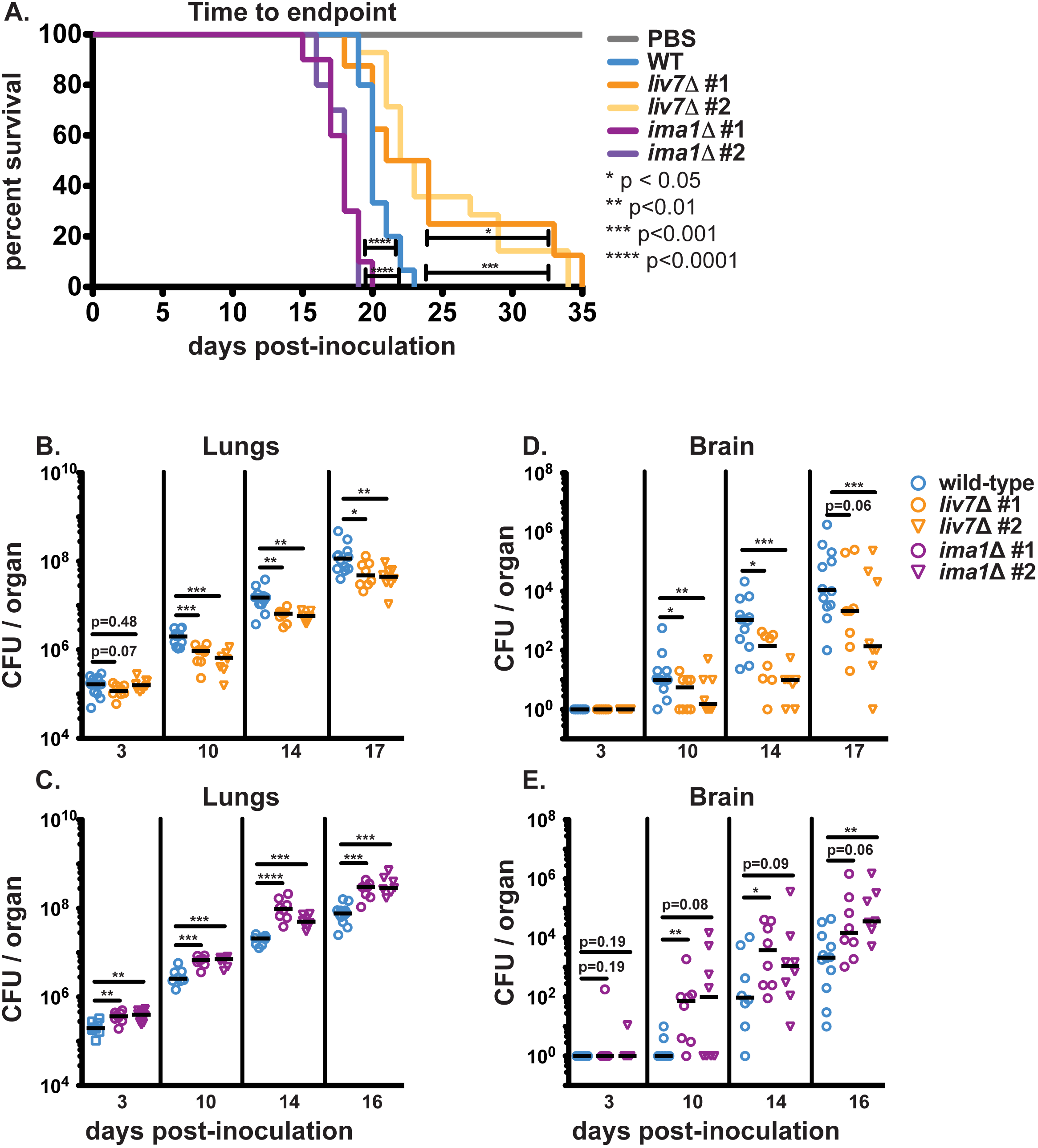
Mutants’ alterations to *in vitro* exo-GXM release correlate with changes in survival and fungal burden during infection. **(A)** C57BL/6NJ mice infected intranasally with *ima1*∆#1 / #2 (n=10 and n=10, respectively) reach endpoint significantly sooner than wild-type infected mice (n=15). Wild-type infected mice reached endpoint sooner than *liv7*∆#1 / #2 (n=8 and n=14, respectively) infected mice. Mock infected animals given sterile 1X PBS (n=5) did not show signs of disease 35 days post-inoculation. P-values were calculated using a Log-rank (Mantel-Cox) Test. **(B and C)** Lung fungal burden is significantly higher in *ima1*∆#1 / #2 (n=8 and n=8, respectively) infected mice than wild-type infected mice (n=8) beginning at least 3 days post-inoculation, while *liv7*∆#1 / #2 infected mice (n=8 and n=8, respectively) show decreased lung burden beginning between 10 days post-inoculation compared to wild-type (n=12). **(D and E)** Dissemination to the brain trends higher in *ima1*∆#1 / #2 infected mice, and is significantly lower in *liv7*∆#1 / #2 infected animals when compared to wild-type P-values were calculated using a Mann-Whitney test.

We also assessed fungal burden by plating homogenized organs for colony forming unit (CFU) counts. Organ fungal burden followed the same trends as survival. Mice inoculated with *liv7*∆#1 / #2 cells consistently presented with lower fungal burden in the lungs by day 10 post-inoculation **(Fig. 5B)**. Dissemination of *liv7*∆#1 / #2 cells to the spleen **(Fig. S3)** and brain **(Fig. 5C)** was also reduced compared to wild-type. In contrast, mice inoculated with *ima1*∆#1 / #2 cells suffered higher pulmonary fungal burden compared to those inoculated with wild-type *C. neoformans* **(Fig. 5D)**. Differences were present as soon as 3 days post-inoculation **(Fig. 5D)**. We also observed a greater number of disseminated *ima1*∆#1 / #2 cells in the liver and spleen throughout the course of infection **(Fig. S3)**. *ima1*∆#1 / #2 cells disseminated to the brain earlier than wild-type cells, with some *ima1*∆#1 / #2 infected mice showing CFUs in the brain as early as 3 dpi **(Fig 5E)**. Total brain fungal burden in *ima1*∆#1 / #2 infected mice trended higher than wild-type, with one independent gene deletion strain achieving a statistically significant increase in fungal burden 10 dpi and beyond, despite high variance in dissemination at the observed time points (**Fig. 5E)**. These results suggest that time-to-endpoint for the mice was at least partially due to fungal lung burden and extrapulmonary dissemination, both of which correlated with *in vitro* exo-GXM secretion.

Since *in vitro* exo-GXM production by *ima1*∆#1 / #2 and *liv7*∆#1 / #2 cells correlated with virulence *in vivo*, we examined whether or not the *in vitro* phenotypes would translate to detectable differences in exo-GXM production in the host environment. We analyzed the levels of GXM in the lungs, livers, spleens and brains of infected mice by performing GXM ELISA’s on 0.22 μm filtered organ homogenates. Exo-GXM levels *in vivo* were highly variable, perhaps reflecting the heterogeneous host environment or assay insensitivity **(Fig. S4)**. In spite of thisvariability, we detected significant reductions in total exo-GXM in the lungs and extrapulmonary organs of *liv7*∆#1 / #2 infected mice at certain time points, with these trends becoming more apparent as infection progressed **(Fig. S4A-D)**. Similarly, *ima1*∆#1 / #2 infected mice displayed increased total exo-GXM in the lungs, spleen and liver by 14 dpi **(Fig. S4E-G)**. We did not observe any interpretable differences in exo-GXM levels on a per cell basis (data not shown), possibly due to changing host conditions over the course of dissemination or assay variability. Spread and/or clearance of exo-GXM within the host likely also played a role, as the spleen and livers of infected mice had massively increased exo-GXM levels on a per cell basis.

Also of note, is that we detected exo-GXM in extrapulmonary organs prior to consistent detection of colony forming units (CFU) in those same organs **(Fig. S5)**. This observation may be relevant for diagnosticians interested in detecting cryptococcal infection prior to dissemination in at-risk patient populations, as early diagnosis of cryptococcosis greatly improves outcomes (48).

### Cell size shifts dramatically during the course of infection parallel to increases in exo-GXM

We investigated whether or not *in vitro* capsule phenotypes for the mutants were recapitulated *in vivo*. We isolated cryptococcal cells from infected mice, fixed them with paraformaldehyde, and measured cell body diameter, cell surface capsule thickness, and total diameter (cell diameter including capsule) using india ink **(Fig. 1A)**.

In wild-type-infected mice, cell and capsule size in the lungs was a broad distribution that shifted significantly over the course of infection, as observed by others (49-51). Large cells were in high abundance early in infection, particularly at 3 dpi **(Fig. 6A)**. These cells were likely Titan cells, which are large, highly polyploid, and increase their size and ploidy through non-mitotic genome replication (14). However, as infection progressed, the frequency of large cells decreased. By 20 dpi, smaller cells around 10μm in total diameter dominated the lungs in number **(Fig. 6A)**. The cell body size and capsule thickness distributions experienced proportional shifts, such that overall cell size to capsule thickness ratios were maintained (**Fig. S6A,B**). In the brain, the distribution of cell and capsule size was much narrower and overlapped with the population of smaller cells in the lungs **(Fig. 6B)**.

**Figure 6:**
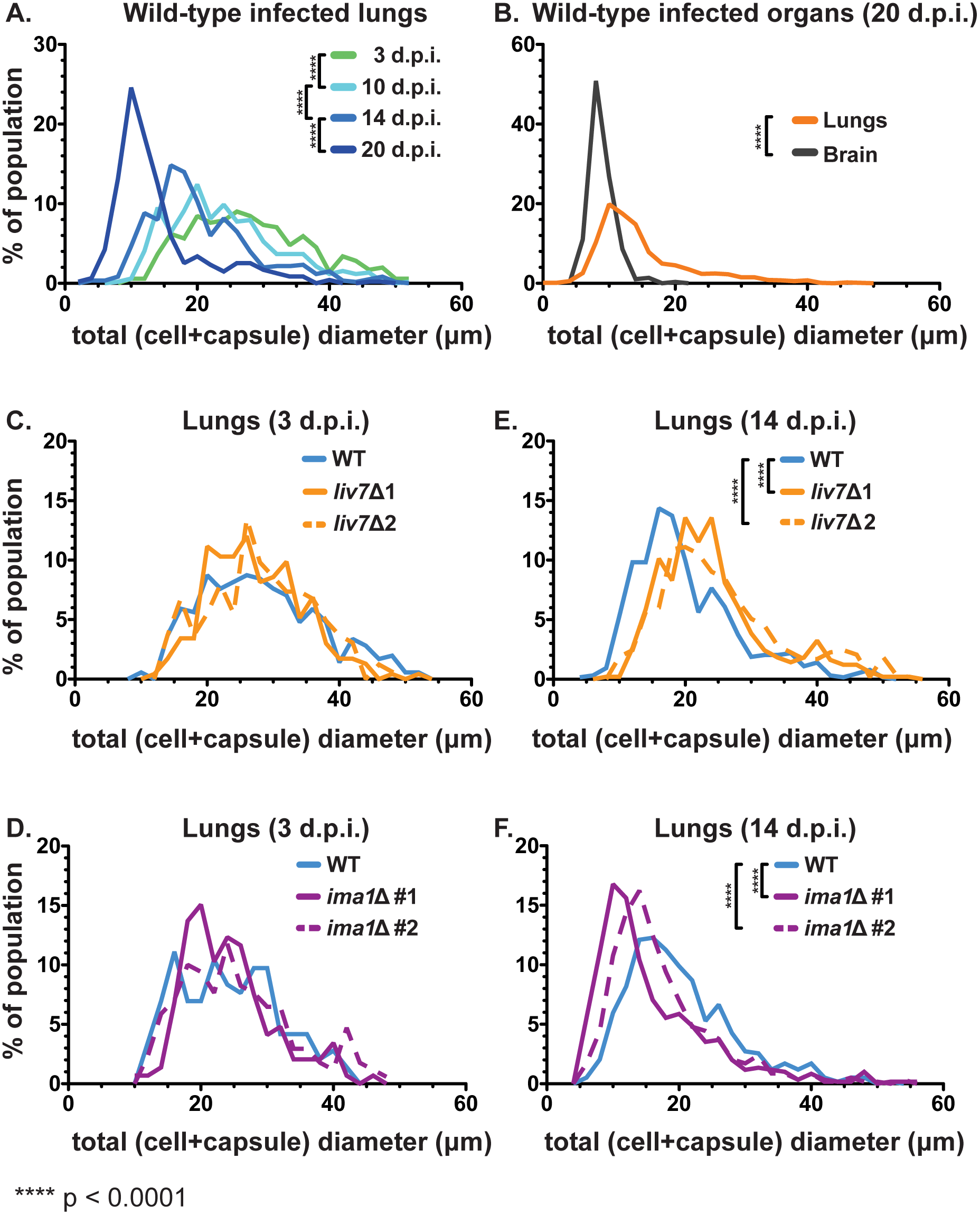
Cell size distributions over the course of infection. We visualized fungal cells from tissue homogenates (from infected mice in **Fig. 5B-E**) in india ink and measured cell surface capsule size. Total diameter = cell + capsule diameter. Cell body diameter = diameter from one edge of the cell wall to the other. Capsule thickness = (total diameter -- cell body diameter)/2 **(Fig 1A)**. **(A)** Mean total cell diameter decreases over time within the lungs of wild-type infected mice as the population shifts toward smaller cells with smaller capsules (n=3-4 mice per time point, ≥120 cells per mouse). **(B)** Disseminated cells found in the brain late in infection (20 dpi) overlay with the size profile of smaller cells found in the lungs at the same time point. **(C and D)** Early after inoculation (3 dpi) the distributions of both **(C)** *liv7*∆#1 / #2 and **(D)** *ima1*∆#1 / #2 cells match that of wild-type in the lungs (n=3 mice, ≥50 cells per mouse). **(E and F)** At an early point in dissemination (14 dpi), **(E)** *liv7*∆#1 / #2 cell populations were of larger average total cell diameter than wild-type *C. neoformans* cells in the lungs. **(F)** *ima1*∆#1 / #2 cells were of smaller average total cell diameter than wild-type *C. neoformans* cells (n=4 mice, ≥120 cells per mouse). P-values were calculated using a Mann-Whitney test.

When we compared the total cell diameter distributions of wild-type and the exo-GXM mutants in the lungs, there was no difference 3 dpi **(Fig. 6C,D)**. By an early time point in dissemination (14 dpi), however, the frequency of smaller cells was higher in *ima1*∆#1 / #2 infected mice and lower in *liv7*∆#1 / #2 infected mice, when compared to wild-type **(Fig. 6E,F)**. The ratio of cell size to capsule thickness was similar amongst all strains **(Fig. S6C).**

Due to this correlation between cell and capsule size and exo-GXM, we hypothesized that levels of exo-GXM could influence cell and capsule size. To test this, we grew cells in strong cell surface capsule-inducing medium (10% Sabouraud’s, pH 7.3) with minimal exo-GXM release. After 24 hours growth at 37°C, we diluted the cultures 1:2 in fresh medium and added 100 ng/ml, 10 μg/ml, or 50 μg/ml of purified GXM. After an additional 24 hours growth, we measured cell and capsule size. We found that both capsule thickness (**Fig. 7A**) and cell size (**Fig. S7A**) decreased in a dosage-dependent manner. The greatest decrease was in capsule thickness, which showed a decrease from a median of 4 μm in control cultures to 1.5 μm in cultures treated with 50 μg/ml GXM, a decrease of 62.5%. 100 ng/ml showed a more modest decrease, to a median capsule thickness of 3.6 μm (a 10% decrease). 50 μg/ml and 10 μg/ml GXM treatments also resulted in a change in cell size, from a median of 6.0 μm for untreated cells to 4.5 μm and 5.3 μm, respectively. 100 ng/ml GXM did not result in a decrease in cell size, despite the observed change in capsule thickness (**Fig. S7A**).

**Figure 7:**
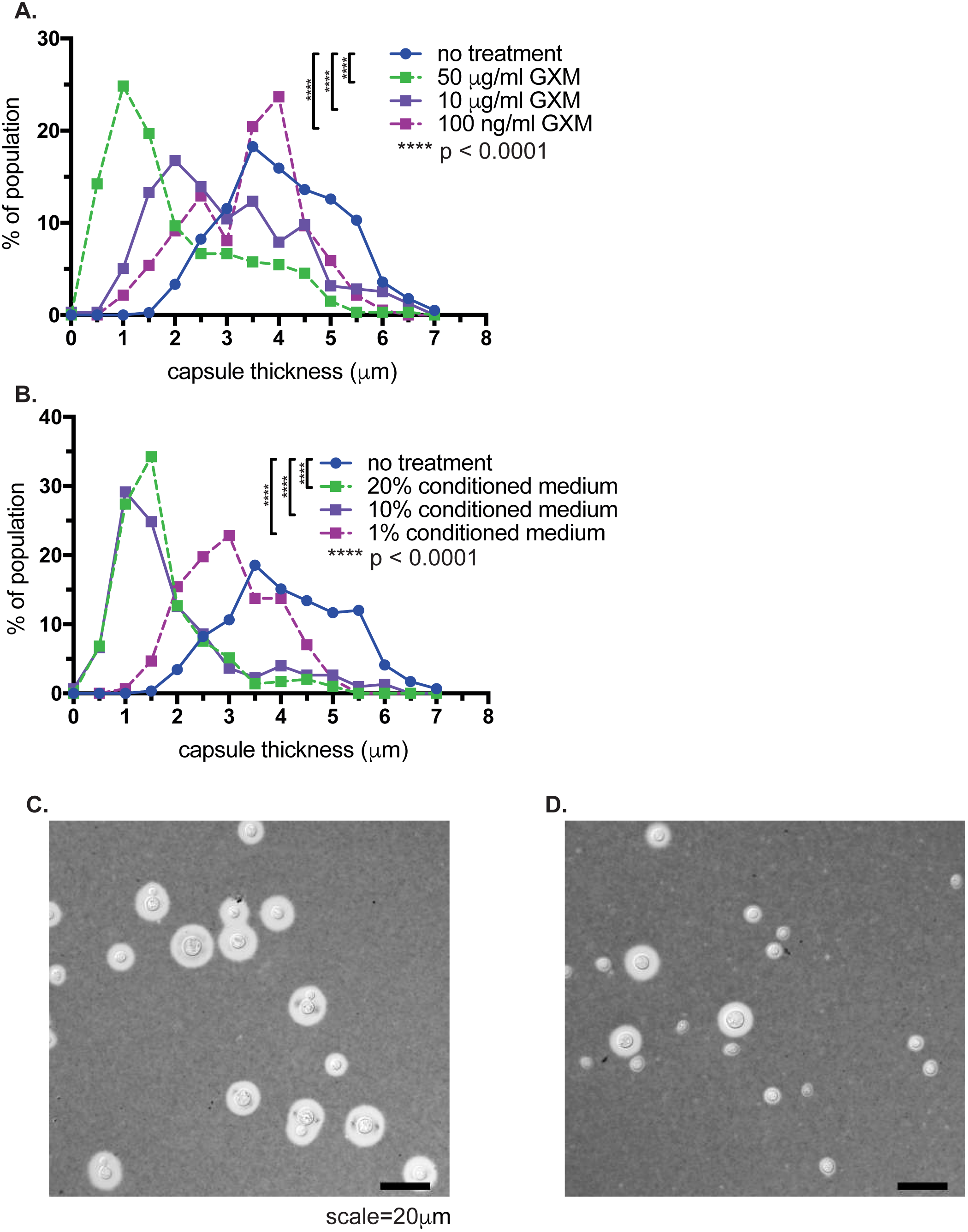
Treatment with GXM decreases capsule thickness. We induced cell surface capsule by growing cells 24 hours in 10% Sabouraud’s pH 7.3, then added various concentrations of either (**A**) purified GXM or (**B**) conditioned medium from a YNB-grown culture of wild-type (KN99) *C. neoformans* cells. We find a dosage-dependent decrease in capsule thickness following exposure to both purified GXM and conditioned medium. Histograms contain data from four separate experiments, with at least 60 cells measured per condition for each experiment. We also observed a decrease in cell size (see **Fig. S7**) with GXM or conditioned medium treatment. P-values were calculated using a Mann-Whitney test. Representative DIC images of **(C)** untreated cells or **(D)** cells treated with 50 μg/ml GXM are shown.

GXM purification can result in contamination by detergents from the purification protocol (30). Thus, we performed the same experiment, but added conditioned medium (from a YNB-grown culture) instead of purified GXM. 20%, 10%, or 1% final concentration of conditioned medium resulted in decreases in both capsule thickness and cell size (**Fig. 7B, Fig. S7**). These capsule and cell size changes also depended on growth: if we did not add fresh medium along with purified GXM, capsule thickness and cell size did not change (**Fig. S7**).

Altogether, these data suggest that changes to exo-GXM observed *in vitro* can affect pathogenesis. Total exo-GXM secreted throughout infection correlated with decreased survival, increased fungal burden and more rapid generation of smaller (haploid) cells in the lungs, which appear more likely to disseminate due to their dominant presence in extrapulmonary organs.

### Exo-GXM limits innate immune cell infiltration into the brain

In human patients, cryptococcal meningoencephalitis is associated with a striking paucity of inflammation (9). The main driver of mortality, particularly in immunocompromised patients, is thought to be excessive fungal burden and GXM accumulation within the CNS, which leads to a devastating increase in intracranial pressure (10). C57BL/6NJ mice infected with the highly virulent KN99 strain display a similar paucity of brain inflammation, despite significant fungal presence. For instance, when we histologically compared the brains of KN99 infected mice to mock-infected animals, we could detect very little sign of infiltrating immune cells by H&E staining in KN99-infected mice, despite local presence of fungi **(Fig. S8)**. This was true both early (14 dpi) **(Fig. S8A,B)** and late (21 dpi) in disseminated infection **(Fig. S8C,D)**.

Considering its immunosuppressive nature, we hypothesized that GXM could very likely play a role in limiting brain inflammation during infection. We correspondingly reasoned that infection with *liv7*∆#1 / #2 cells might result in increased immune infiltration into the brain, due to *liv7*∆#1 / #2 cell’s reduced exo-GXM secretion.

In order to address this hypothesis, we harvested the brains of wild-type and *liv7*∆#1 / #2 infected animals at 20 days post-intranasal inoculation and analyzed immune infiltration into the brain via flow cytometry. CD4+ **(Fig. 8A)** and CD8+ **(Fig. 8B)** cells were scarce in both wild-type and *liv7*∆#1 / #2-infected brains. These data suggest that T cells do not significantly respond to brain invasion by *C. neoformans*. Innate immune cells (macrophages/neutrophils) did show some response to wild-type *C. neoformans* cells in the brain, but it was only slightly elevated when compared to mock-infected animals **(Fig. 8C,D)**. This is in stark contrast to bacterial or viral meningitis, which often show high levels of infiltrating neutrophils and macrophages (7, 8). Infiltration of both macrophages and neutrophils was increased in *liv7*∆#1 / #2 infected brains **(Fig. 8C,D)**. These results suggest that exo-GXM likely plays an important role in brain immunosuppression that is independent of surface capsule.

**Figure 8:**
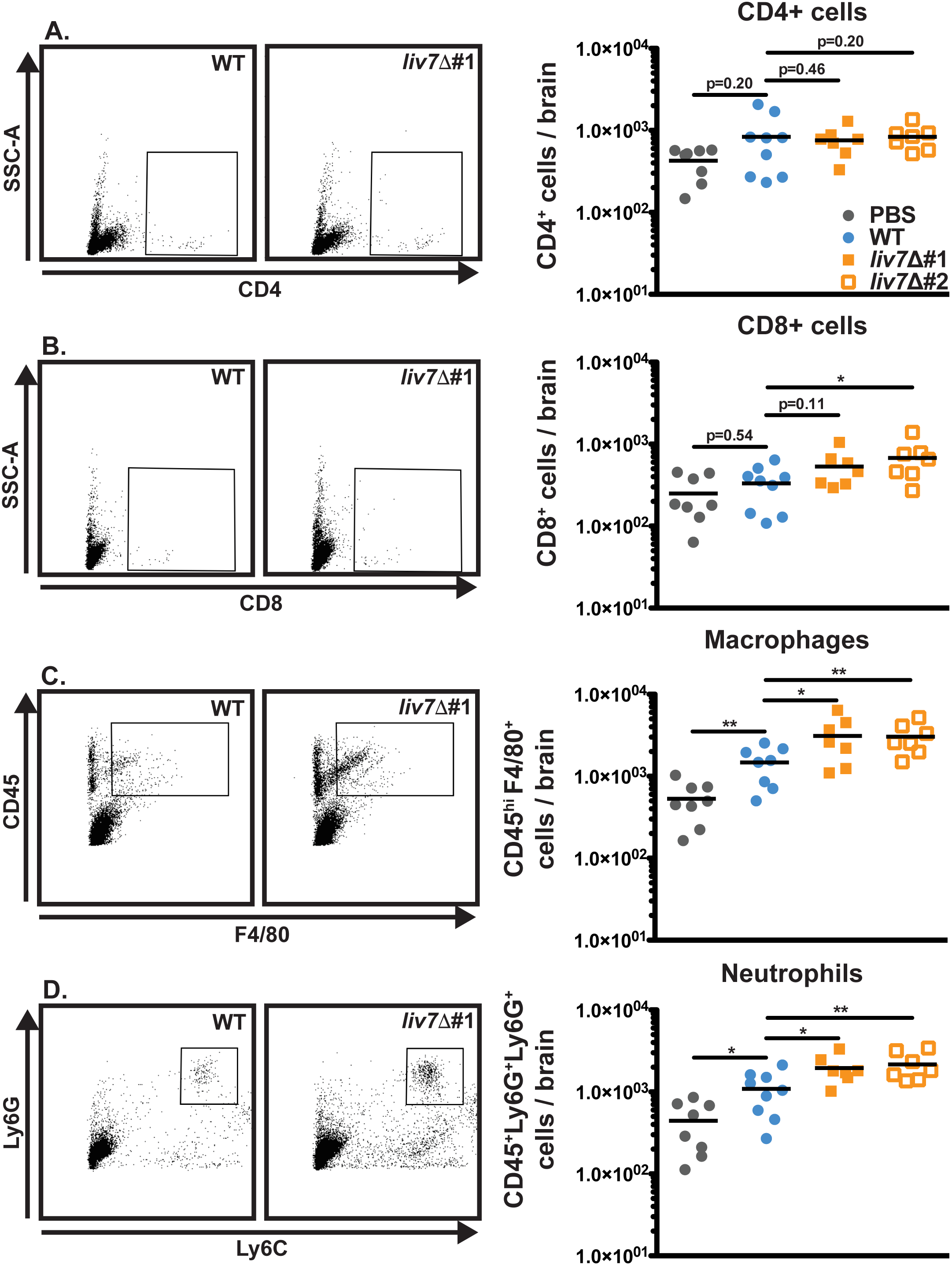
Mice infected with *liv7*∆ cells display increased innate immune infiltrate in the brain. Mouse brains were harvested late (20 dpi) in infection for flow cytometry analysis of infiltrating immune cells. **(A)** CD4^+^ T cells are scarce in both wild-type and *liv7*∆#1 / #2 infected brains. **(B)** CD8^+^ T cells show a significant increase over wild-type in *liv7*∆#2 infected brains, but this was not replicated in *liv7*∆#1 infected brains **(C)** Macrophages (CD45^hi^F4/80^+^) and **(D)** neutrophils (CD45^+^Ly6G^+^Ly6C^+^) are significantly increased in the brains of *liv7*∆#1 and #2 as compared to wild-type and mock-infected brains. P-values were calculated using a Mann-Whitney test; bars represent the median.

We next sought to determine if exo-GXM was sufficient to suppress immune infiltration into the brain if we induced brain inflammation by direct intracranial inoculation. We purified GXM from YNB-grown cultures using standard methods (30). Since we detected GXM associated with the brain up to five days prior to the appearance of CFU **(Fig. S5)**, we administered 200 μg of purified GXM daily by intraperitoneal injection, beginning five days prior to inoculation **(Fig. 9A)**. Additional mice were administered sterile PBS as a control. We then inoculated mice intracranially with either wild-type KN99 or acapsular *cap60*∆ cells. Unsurprisingly, *cap60*∆ cells elicited greater numbers of immune infiltration into the brain **(Fig. 9B,C)**, and achieved a significantly lower fungal burden than wild-type *C. neoformans* **(Fig. 9D)**. However, administration of GXM to mice infected with *cap60*∆ cells reduced immune infiltration (CD45^hi^ cells) into the brain **(Fig. 9B,C and Fig. S9)**, and increased fungal burden when compared to PBS-treated mice **(Fig. 9D)**. These results demonstrate that in the context of an inflammatory infection, exo-GXM is sufficient to promote fungal survival in the brain, likely through the suppression of brain immune infiltration.

**Figure 9:**
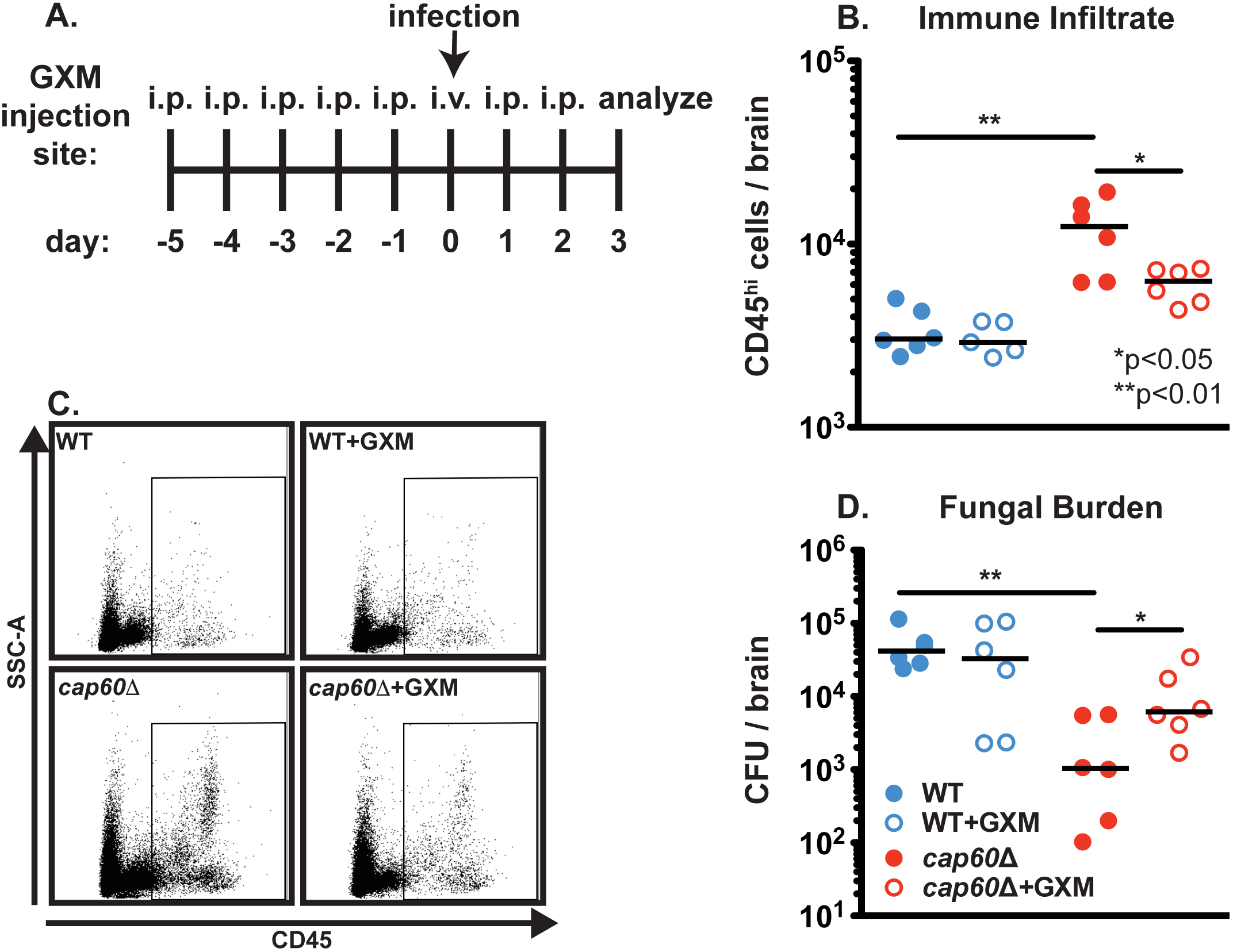
Purified GXM is sufficient to suppress immune infiltration into the brain in response to inflammation-inducing acapsular (*cap60*∆) *C. neoformans*. 6-week-old C57Bl/6NJ mice were intracranially inoculated with 200 *cap60*∆ fungal cells in 30 μl 1X PBS. Beginning 5 days prior to inoculation, mice were administered intraperitoneal injections of either 200 μg/mL GXM or 200 μl sterile PBS. On the day of inoculation, mice were administered this treatment intravenously to ensure GXM would be present in the blood-stream. At 3 dpi brains were harvested to determine fungal burden by colony forming unit counts. Separate mice were sacrificed to analyze infiltrating immune cells by flow cytometry. **(A)** Diagram of experimental procedures. **(B)** Mice infected with *cap60*∆ displayed increased brain immune infiltrate (CD45^hi^ cells) over wild-type infected mice. Immune infiltration into the brains of *cap60*∆ infected mice was reduced with the administration of GXM. **(C)** Representative flow plots for data shown in **(B)**. **(D)** Mice infected with wild-type KN99 cells suffered increased fungal brain burden as compared to mice infected with *cap60*∆. Administration of GXM had no significant effect on wild-type fungal burden, but resulted in significantly increased *cap60*∆ fungal burden compared to *cap60*∆-infected mice that did not receive GXM. P-values were calculated using a Mann-Whitney test; bars represent the median.

## Discussion

Surface capsule is critical for *C. neoformans* virulence. However, GXM that is not attached to the cell surface, or exo-GXM, accumulates to significant levels in laboratory culture and during infection (10, 30, 31). Our data strongly suggest that *C. neoformans* inversely regulates surface capsule formation and exo-GXM release according to environmental cues. Within our tested conditions, GXM was constitutively produced but alternately retained at the cell surface or released into the extracellular milieu. Previous findings have also indicated that exo-GXM release might be an active process. For instance, a study comparing the properties of exo-GXM and capsular GXM showed that despite sugar composition remaining the same, capsular GXM and exo-GXM manifested distinct biophysical and antigenic properties (34). Additionally, electromobility of exo-GXM decreases under capsule inducing conditions, implying that structural changes that influence capsule formation (35). We also observed that *O*-acetylation of GXM’s mannose backbone changes with environmental conditions. These findings potentially suggest that differential regulation of surface capsule and exo-GXM could occur at the level of GXM polymer length and/or other modification. More work is required to elucidate biophysical differences between cell surface retained- and exo-GXM.

We identified genes that play a role in exo-GXM release. Deletion of *LIV7* reduces exo-GXM release in rich growth medium when cell surface capsule does not form, but does not affect capsule thickness. It has been previously demonstrated that *LIV7* is important for virulence and likely functions in Golgi transport (42, 44). Our second exo-GXM mutant, a deletion of the gene *IMA1*, increased exo-GXM release under strong capsule inducing conditions without affecting capsule thickness. We used these two exo-GXM mutants as tools to investigate the biological importance of exo-GXM independent of surface capsule.

We first established a positive correlation of exo-GXM release with biofilm adherence, suggesting that exo-GXM release during environmental growth may be important for promoting community level structure and adherence. It would not be surprising for there to be additional functions for exo-GXM in environmental settings.

In a murine infection model, we showed a correlation between elevated *in vitro* exo-GXM levels, fungal burden and poor host survival. Other groups have also connected varied exo-GXM release with changes to virulence. Analysis of a virulence-associated transcriptional network map previously revealed a positive correlation with exo-GXM release and mouse lung infectivity over 7 days (52). However, the transcription factor mutants also had altered surface capsule thickness, which may have influenced infectivity (52). Deletion of the flippase encoding gene *APT1* also resulted in reduced *in vitro* exo-GXM release despite wild-type surface capsule. The knockout was hypovirulent, but in contrast to our mutants, had reduced surface capsule thickness *in vivo* (53). Our results support these previous findings, and our new exo-GXM mutants are a powerful tool for investigating exo-GXM because they do not suffer any alterations to additional virulence factors. Our data also provide additional support for a model in which regulated release of exo-GXM enhances virulence independent of surface capsule.

Interestingly, exo-GXM also correlated with changes in cell body and capsule size distributions in the lungs. In wild-type *C. neoformans* infected animals, fungal cell body size and capsule thickness decreased over the course of infection, as exo-GXM levels increased. Correspondingly, increased GXM levels in the mouse lungs positively correlated with an increased frequency of smaller cells at an early time point in dissemination. *C. neoformans* cells in the brain and other extrapulmonary organs are much smaller than the lungs (**Fig. 6B** and (50, 54, 55)), suggesting that the emergence of smaller cells in the lungs is an important step in dissemination. The addition of purified GXM to *C. neoformans* cells growing in capsule-inducing media was sufficient to decrease cell body size and capsule thickness in a growth-dependent manner (**Fig. 7**). These data suggest that exo-GXM may actually provide a concentration-dependent signal to *C. neoformans* cells that reduces cell size and capsule thickness. In the lungs, this mechanism may be a contributing factor in the generation of small cells with a greater propensity for dissemination.

There is large body of literature demonstrating immunosuppressive properties for GXM (19). We focused on the brain, as cryptococcal meningoencephalitis is the leading cause of death in cryptococcosis patients and is characterized by low levels of inflammation (9). Here, we observed that deleting a gene required for wild-type levels of exo-GXM release *in vitro* (*LIV7*) altered the host immune response to *C. neoformans* brain infection. Mice infected with *liv7*∆ cells had increased macrophages and neutrophils infiltrating the brain, compared to wild-type infected mice. Furthermore, administration of purified GXM was sufficient to reduce brain infiltrating immune cells in the context of acapsular *C. neoformans* infection. These data echo a previous study that showed GXM could reduce early infiltration of neutrophils in a model of acute bacterial meningitis (56). Our results suggest that exo-GXM is an actively secreted virulence factor that may influence cell morphology to facilitate dissemination, and is capable of distally suppressing immune infiltration into the brain.

**Table S1:** Exo-GXM mutant screen results. We screened the *C. neoformans* partial knockout collection (CM18 background, 1200 targeted gene knockouts) under YNB, which results in high exo-GXM release by wild-type cells, or 10% Sabouraud’s pH7.3, which results in low exo-GXM release. **(A)** Gene deletions which resulted in reduced exo-GXM release in YNB after 24 hours but no growth defect or a substantial reduction (>25%) reduction in cell surface capsule thickness in 10% Sabouraud’s pH 7.3. **(B)** Gene deletions which resulted in increased exo-GXM release in 10% Sabouraud’s pH 7.3 after 24 hours. Class 1 gene deletion mutants had approximately wild-type-sized capsule thickness, while Class 2 mutants had reduced capsule thickness in 10% Sabouraud’s pH 7.3.

| **(A) Gene deletion mutants with reduced exo-GXM in YNB** |  |  |
| --- | --- | --- |
| **CNAG** | **Molecular Function** | **Capsule** |
| CNAG_03171 | WEE protein kinase | WT (++) |
| CNAG_04461 | DNA helicase | ++ |
| CNAG_06673 | conserved hypothetical protein | ++ |
| CNAG_04262 | E3 ubiquitin-protein ligase (NRDP1) | ++ |
| CNAG_04443 | conserved hypothetical protein | ++ |

| **(B) Gene deletion mutants with increased exo-GXM in 10 Sabouraud's, pH 7.3** |  |  |  |  |  |
| --- | --- | --- | --- | --- | --- |
| **CNAG** | **Mutant Class** | **Molecular Function** | **Capsule** | **Urease** | **L-DOPA** |
| CNAG_00658 | 1 | hypothetical protein | WT (++) | WT (++) | WT (++) |
| CNAG_03188 | 1 | histone-lysine N-methyltransferase SETD2 | ++ | ++ | -/+ |
| CNAG_05838 | 1 | Rho GTPase activating protein | +++ | ++ | +++ |
| CNAG_02189 | 1 | alpha-amylase | ++ | ++ | ++ |
| CNAG_04756 | 1 | hypothetical protein | ++ | ++ | ++ |
| CNAG_01551 | 2 | transcription factor (Gat201) | -/+ | ++ | ++ |
| CNAG_03582 | 2 | pH response regulator protein (Rim20) | -/+ | +++ | -/+ |
| CNAG_00375 | 2 | histone acetyl transferase | -/+ | ++ | -/+ |
| CNAG_05690 | 2 | histone deacetylase (Rpd3) | -/+ | ++ | ++ |
| CNAG_04863 | 2 | ESCRT-II complex subunit (Vps25) | -/+ | +++ | -/+ |
| CNAG_03202 | 2 | adenylate cyclase | - | ++ | -/+ |
| CNAG_00761 | 2 | coiled-coil domain-containing protein | + | +++ | -/+ |
| CNAG_06606 | 2 | Rho family protein | + | +++ | + |
| CNAG_05901 | 2 | hypothetical protein | + | + | -/+ |
| CNAG_02215 | 2 | transcriptional activator (Hap3) | + | + | -/+ |

**Figure S1:**
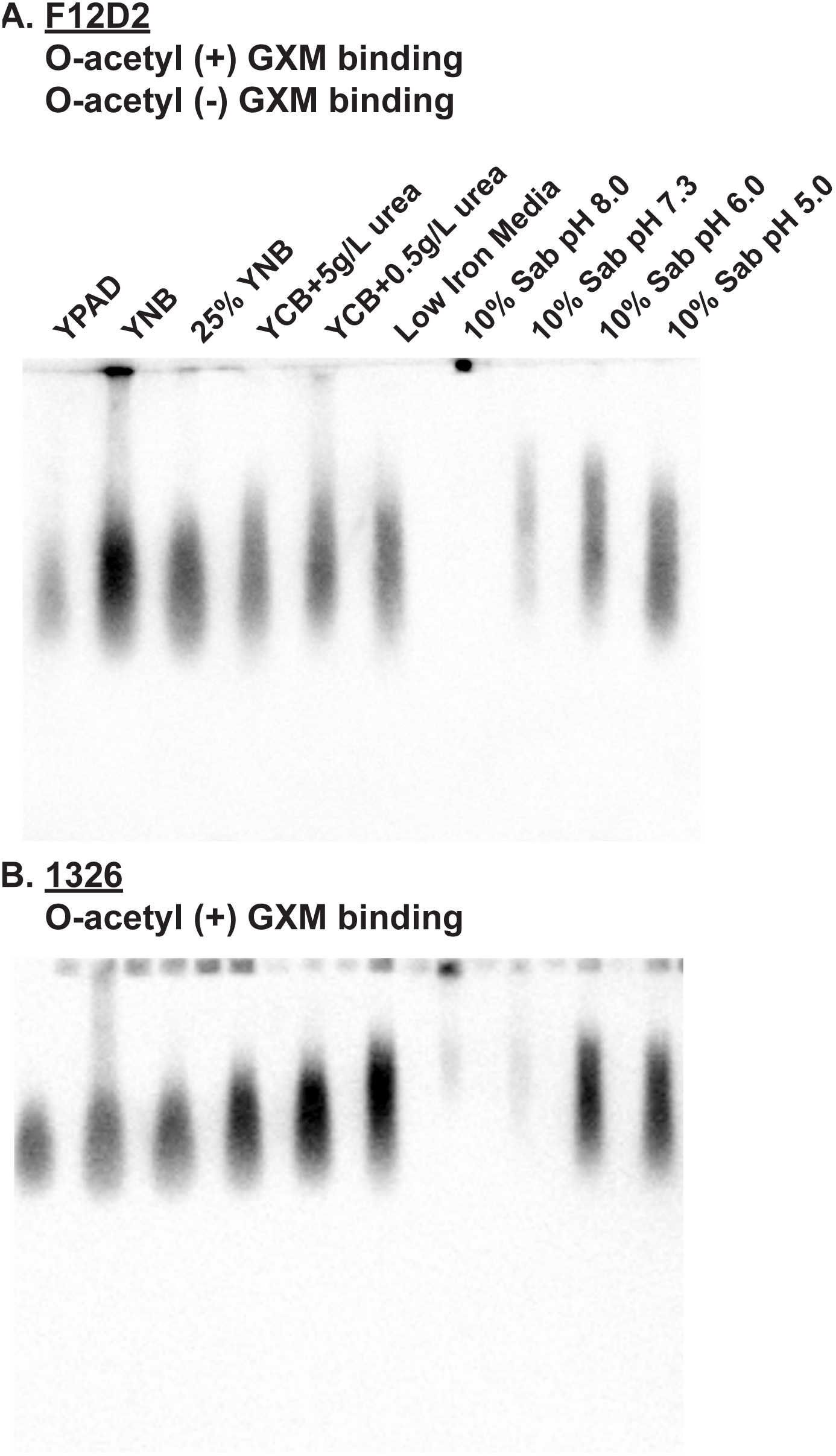
Proportion of *O*-acetylated exo-GXM increases under stronger capsule inducing conditions. Conditioned media was collected and blotted as in **Fig. 1**. **(A)** Detection of GXM with an acetylation insensitive mAb (F12D2). **(B)** Detection of GXM from the same conditioned media with an acetylation-sensitive mAb (1326), which only recognizes *O*-acetylated GXM. Increased intensity indicates a greater level of *O-*acetylated GXM.

**Figure S2:**
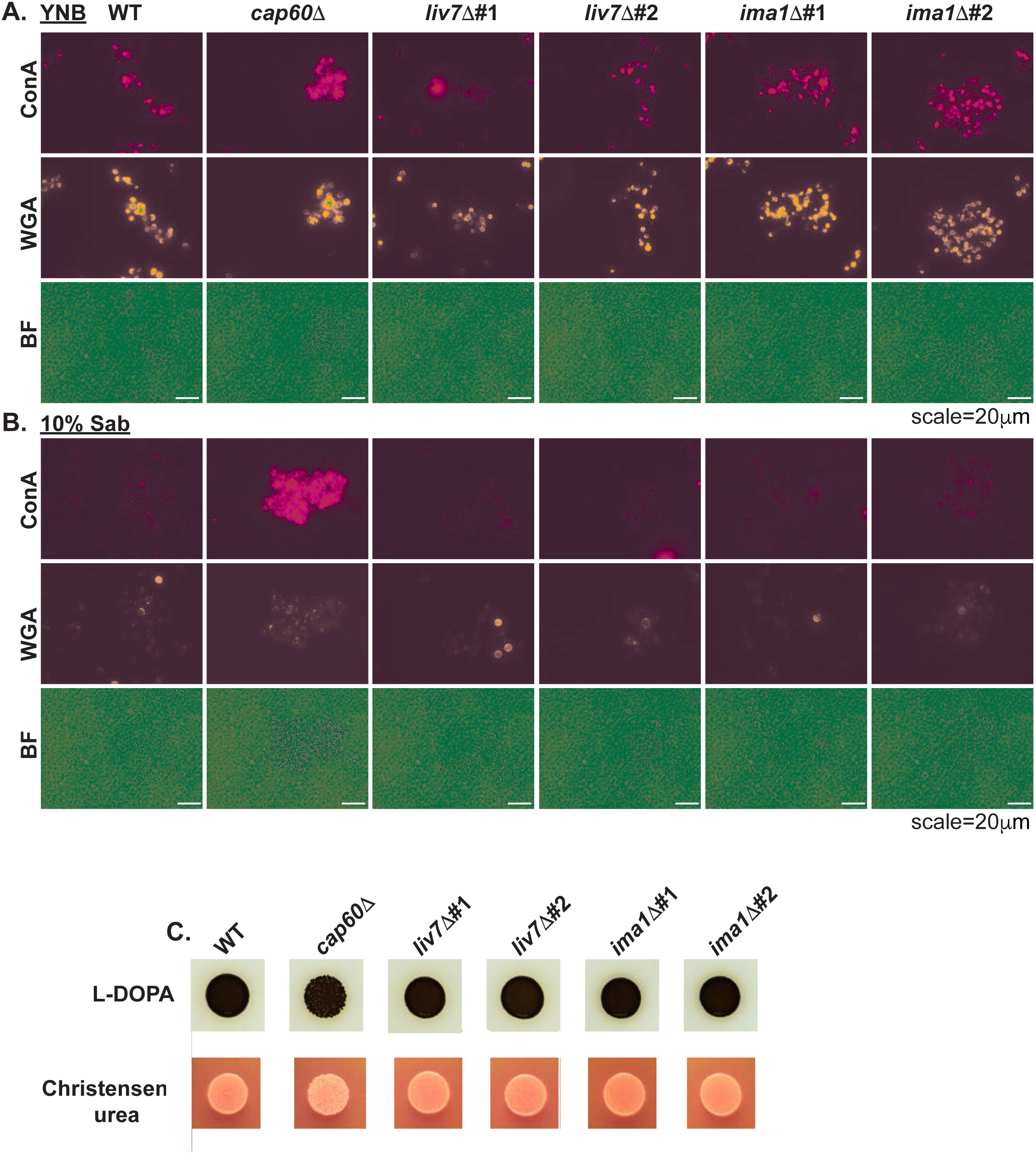
Canonical virulence determinants are intact in *liv7*∆ and *ima1*∆ cells. **(A)** Cells were grown overnight in YNB + 2%glucose, stained with the fluorescently labeled lectins concanavalin A (ConA-rhodamine) and wheat germ agglutinin (WGA-fluorescein) to estimate exposure of PAMPs on the cell surface. **(B)** Cells were grown overnight in YNB + 2%glucose, subcultured 1:100 in 10% Sabouraud’s dextrose (10% sab), pH 7.3, and stained as in **(A)**. PAMP exposure was similar across all strains except *cap60*∆, which lacks surface capsule. **(C)** 2.5x10^4^ cells were spotted on L-3,4-dihydroxyphenylalanine (L-DOPA) agar to observe melanization 48 hours later. No obvious differences were detected. **(D)** 2.5x10^4^ cells were spotted on Christensen’s urea agar to observe urease secretion 48 hours later as the change in agar coloration from orange to pink. No obvious differences were detected.

**Figure S3:**
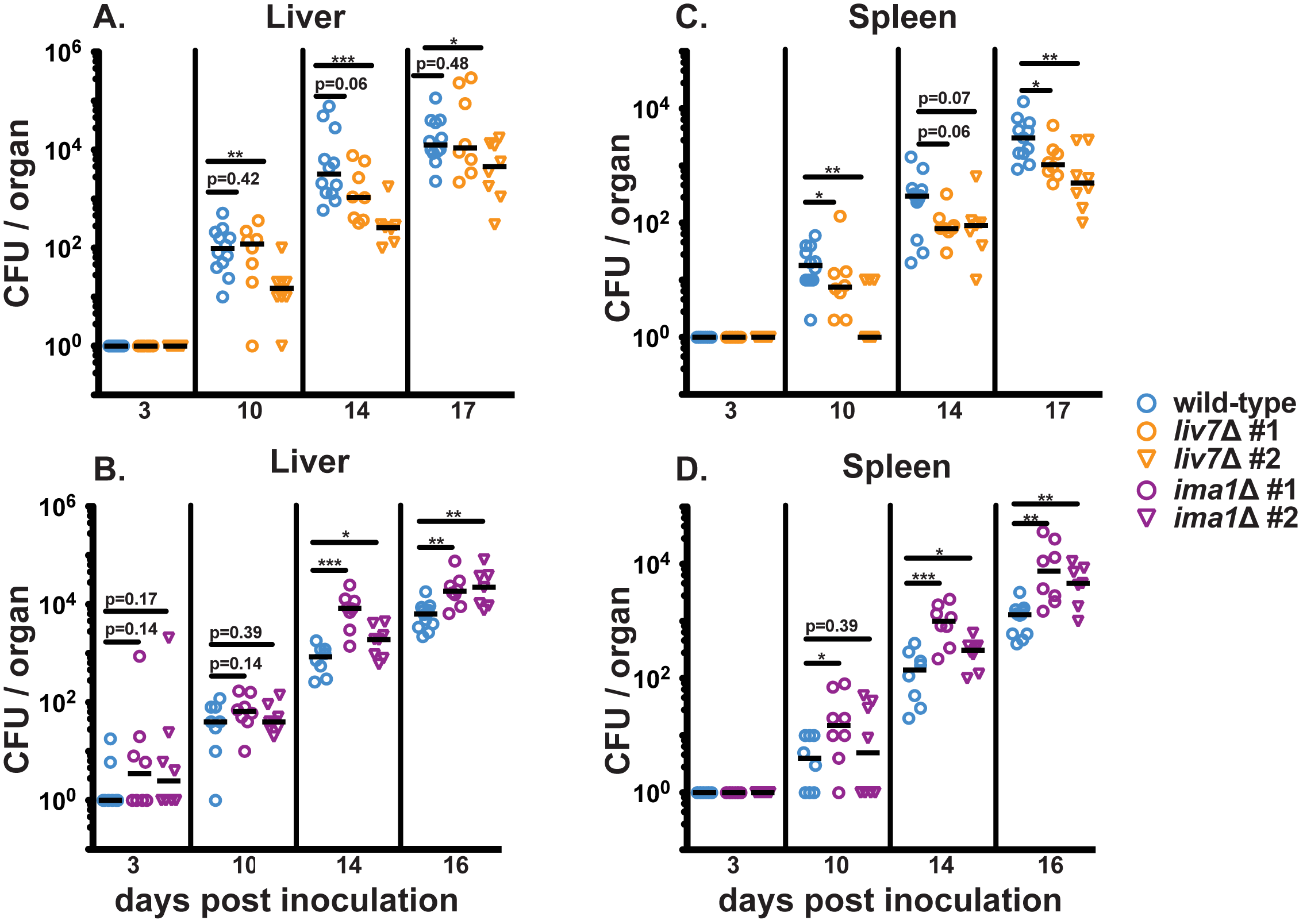
Liver and spleen fungal burden mostly correlates with *in vitro* exo-GXM production. These data are from the same experiments as in **Fig. 5**. **(A)** Fungal burden in the livers of wild-type and *liv7*∆#1 / #2 infected mice did not show consistent differences over the course of infection. **(B)** Fungal burdens in *ima1*∆#1 / #2 infected livers were significantly higher than wild-type at 14 and 16 dpi. **(C)** Fungal burden in the spleens of *liv7*∆#1 / #2 infected mice was significantly lower than wild-type-infected mice at 10 and 17 dpi. **(D)** Fungal burdens in *ima1*∆#1 / #2-infected spleens were significantly higher than wild-type infected mice at 14 and 16 dpi. P-values were calculated using a Mann-Whitney test.

**Figure S4:**
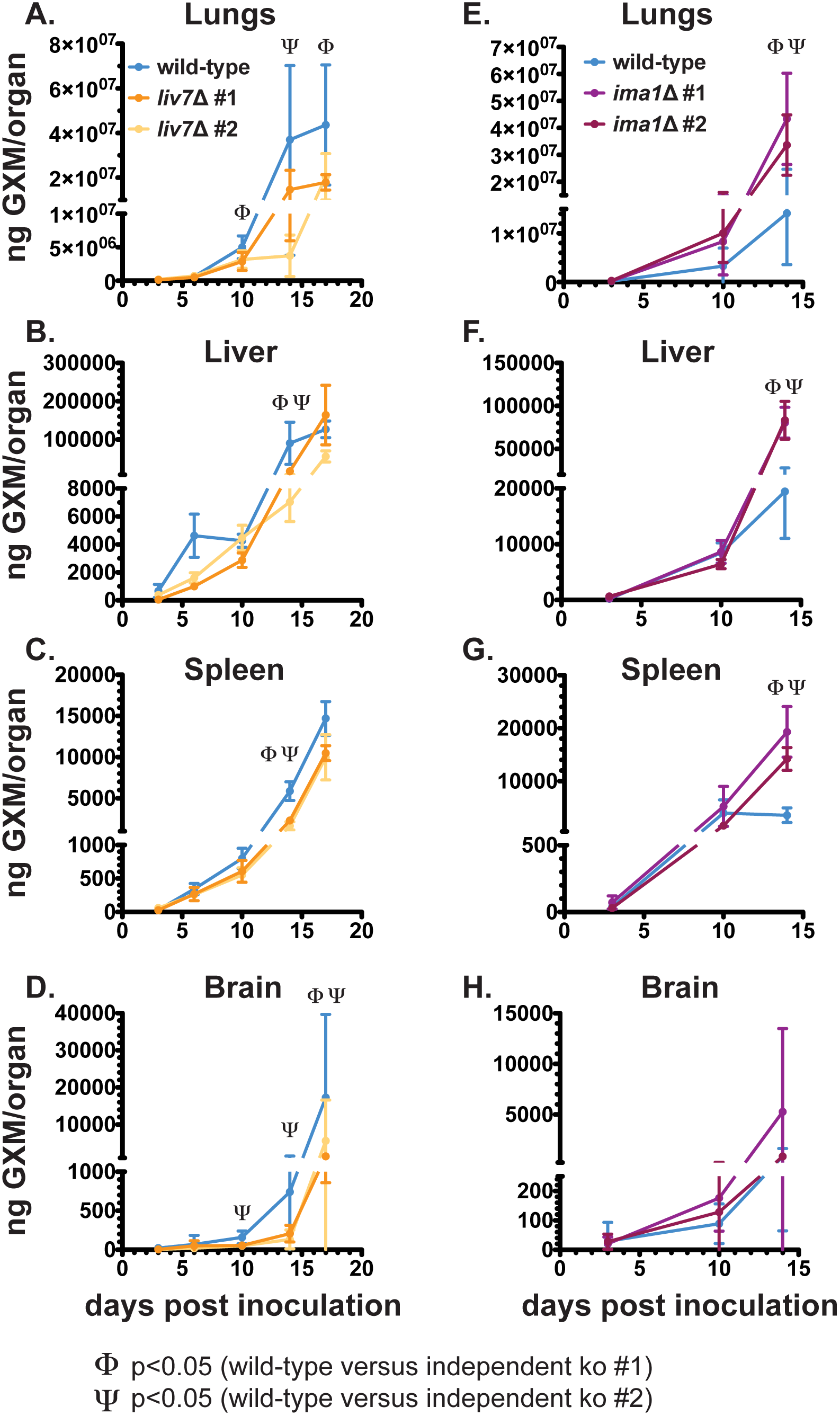
Total free GXM levels in mice infected with *C. neoformans* exo-GXM mutants. Tissues from mice in **Fig. 3B-G** were homogenized and passed through a 0.22 μm filter to remove fungal cells, then GXM levels were measured by ELISA. **(A)** Mouse lungs infected with *liv7*∆#1 / #2 cells showed trends toward reductions in exo-GXM levels when compared to wild-type, though statistical significance is not consistent across independent gene deletions. **(B and C)** Mouse livers and spleens infected with *liv7*∆#1 / #2 cells showed reduced exo-GXM levels when compared to wild-type-infected organs at 14 dpi. **(D)** Mouse brains infected *liv7*∆#1 / #2 cells showed reduced exo-GXM when compared to wild-type at 17dpi. **(E-G)** Exo-GXM was increased in *ima1*∆#1 / #2 infected lungs, livers, and spleens when compared to wild-type-infected organs at 14dpi. **(H)** No significant differences in exo-GXM were observed in *ima1*∆#1 / #2 infected brains when compared to wild-type-infected brains. P-values were calculated using a Mann Whitney t test.

**Figure S5:**
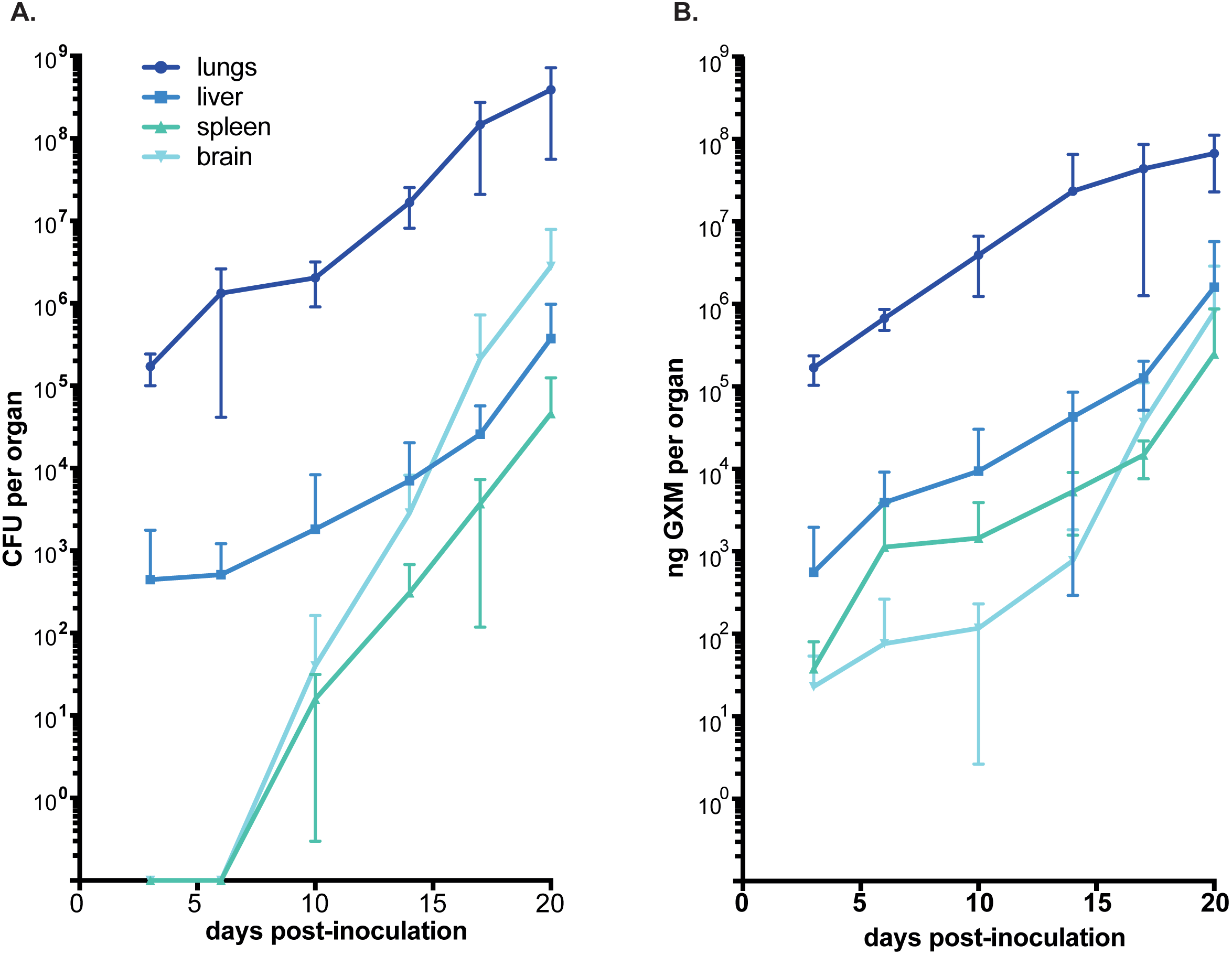
GXM appears in brains and spleens prior to the appearance of CFU. A time course of **(A)** fungal burden (CFU) and **(B)** GXM per organ following infection with wild-type *C. neoformans* shows that GXM is detectable in all organs by 3 dpi. CFU were not detectable in brains or spleens until 10 dpi. These data are the compiled wild-type infection data from **Fig. 5**, **Fig. S3**, and **Fig. S4**.

**Figure S6:**
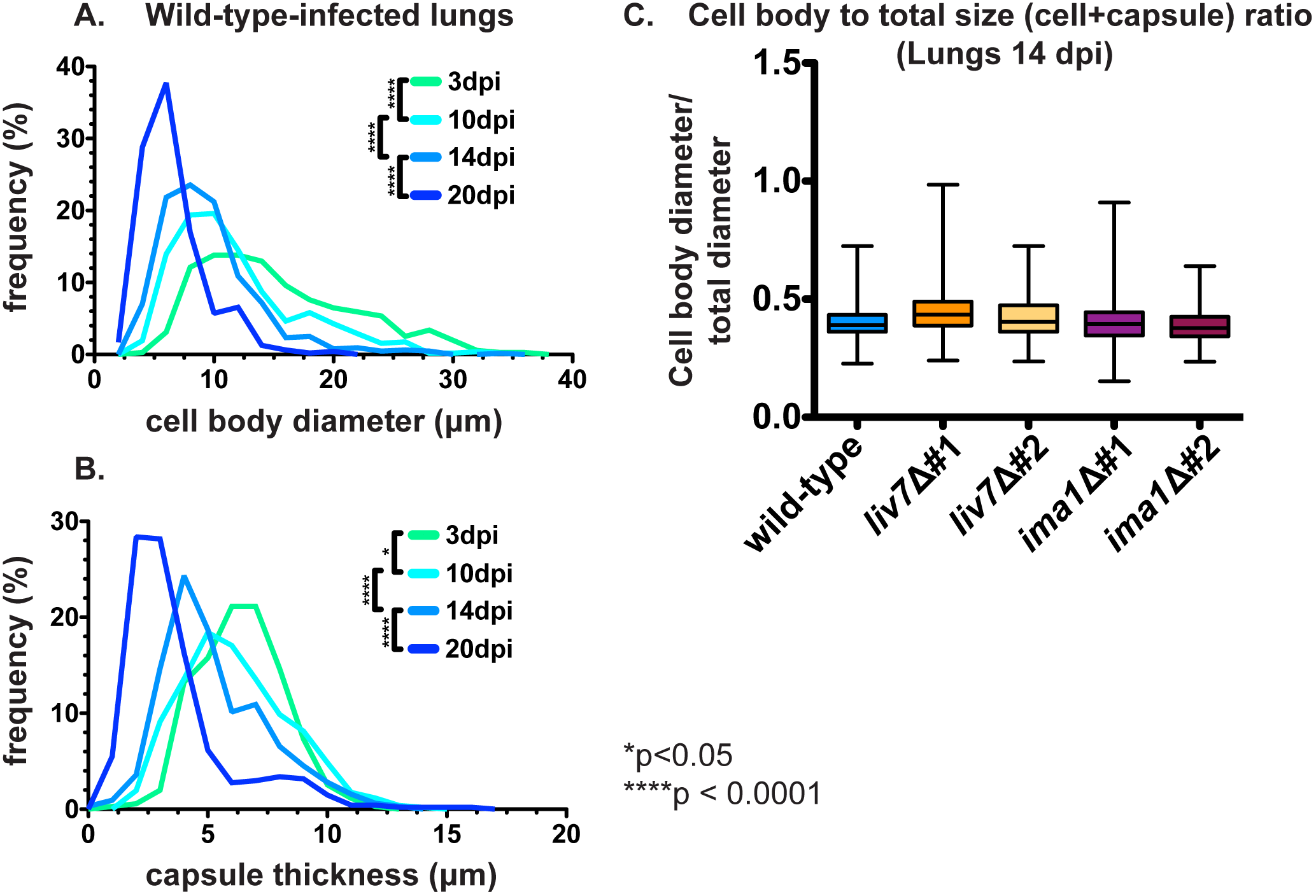
Distribution of *C. neoformans* cell body diameter and cell capsule thickness shift over the course of lung infection: These data are from the same experiments as **Fig. 6**. **(A)** Average *C. neoformans* cell body diameter in the lungs decreases over the course of infection (n=3-4 mice per time point, ≥120 cells per mouse). **(B)** Average capsule thickness in the lungs decreases over the course of infection at a rate similar to the change in cell body diameter (n=3-4 mice per time point, ≥120 cells per mouse). **(C)** The proportion of cell size to capsule thickness in the lungs is similar across wild-type, *liv7*∆#1 / #2, and *ima1*∆#1 / #2 cells in the lungs (n=4 mice, ≥120 cells per mouse). P-values were calculated using a Mann-Whitney test; error bars show medians.

**Figure S7:**
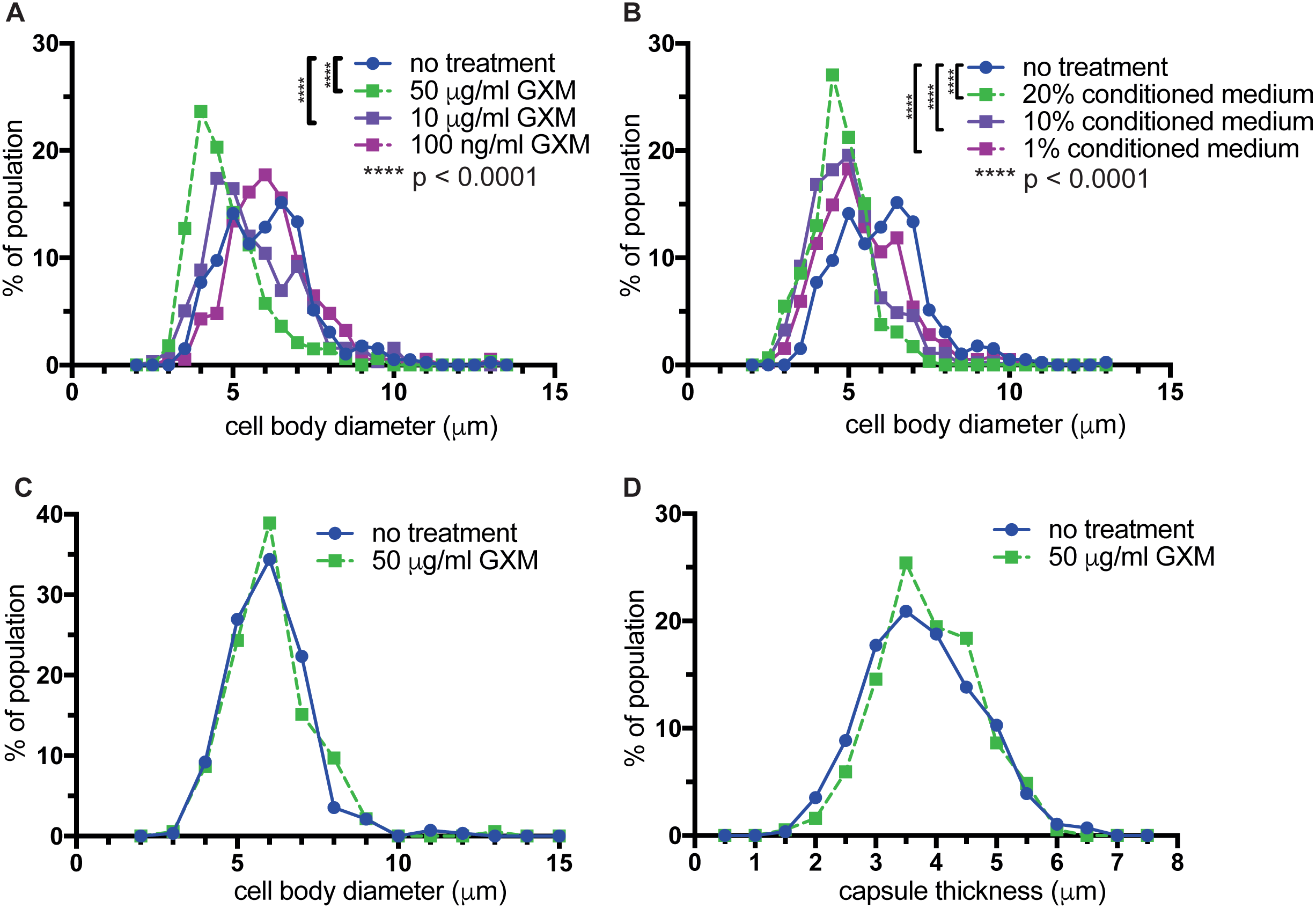
Treatment with GXM decreases cell size. These data are from the same experiments as **Fig. 7**. Cell size decreases in a dosage-dependent manner with the addition of (**A**) purified GXM at 50 μg/ml or 10 μg/ml, but not 100 ng/ml, even though capsule thickness decreased with the addition of 100 ng/ml GXM. (**B**) Conditioned medium at final concentrations of 20%, 10%, or 1% all decrease cell size. (**C**) Cell size and (**D**) capsule thickness do not change if cultures are not administered additional growth medium (10% Sabouraud’s, pH 7.3) along with purified GXM, suggesting that these size changes are growth-dependent. P-values were calculated using a Mann-Whitney test.

**Figure S8:**
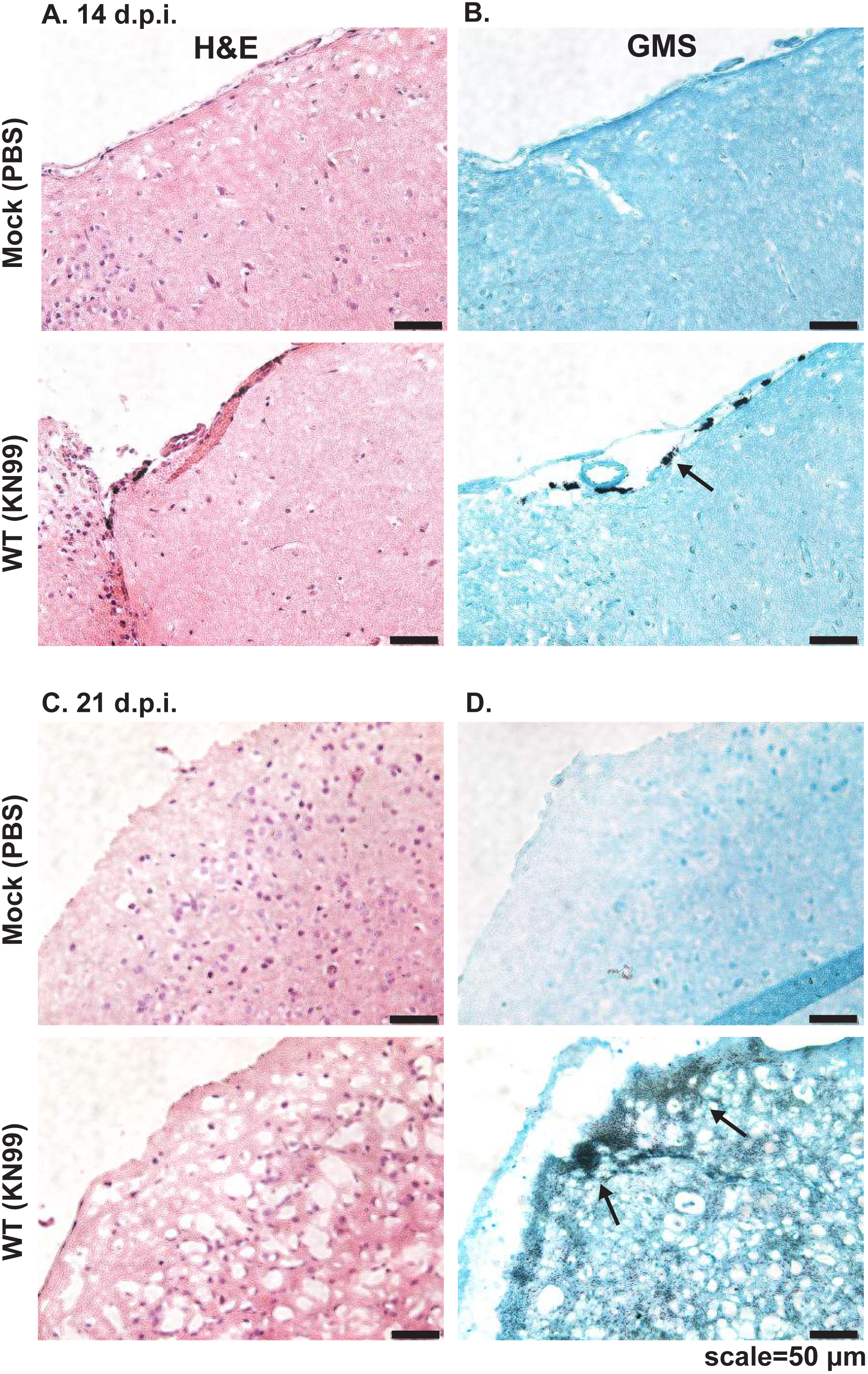
Few immune cells infiltrate the brains of mice with disseminated cryptococcosis, despite high fungal burden. **(A)** Representative hematoxylin and eosin (H&E) and **(B)** consecutive Grocott’s methenamine silver (GMS) stained midbrain sections early (14 dpi) in brain infection. We observed no signs of inflammatory infiltrate (excess purple hematoxylin staining) and minimal fungal presence (black silver staining; arrows point to fungi) early. **(C)** Representative H&E and **(D)** GMS stained cerebral cortex sections late (21 dpi) in brain infection. We continued to detect few signs of inflammatory infiltrate in H&E stained sections late in infection, despite significant and diffuse fungal presence within the meninges and parenchyma of the brain (arrows point to fungi).

**Figure S9:**
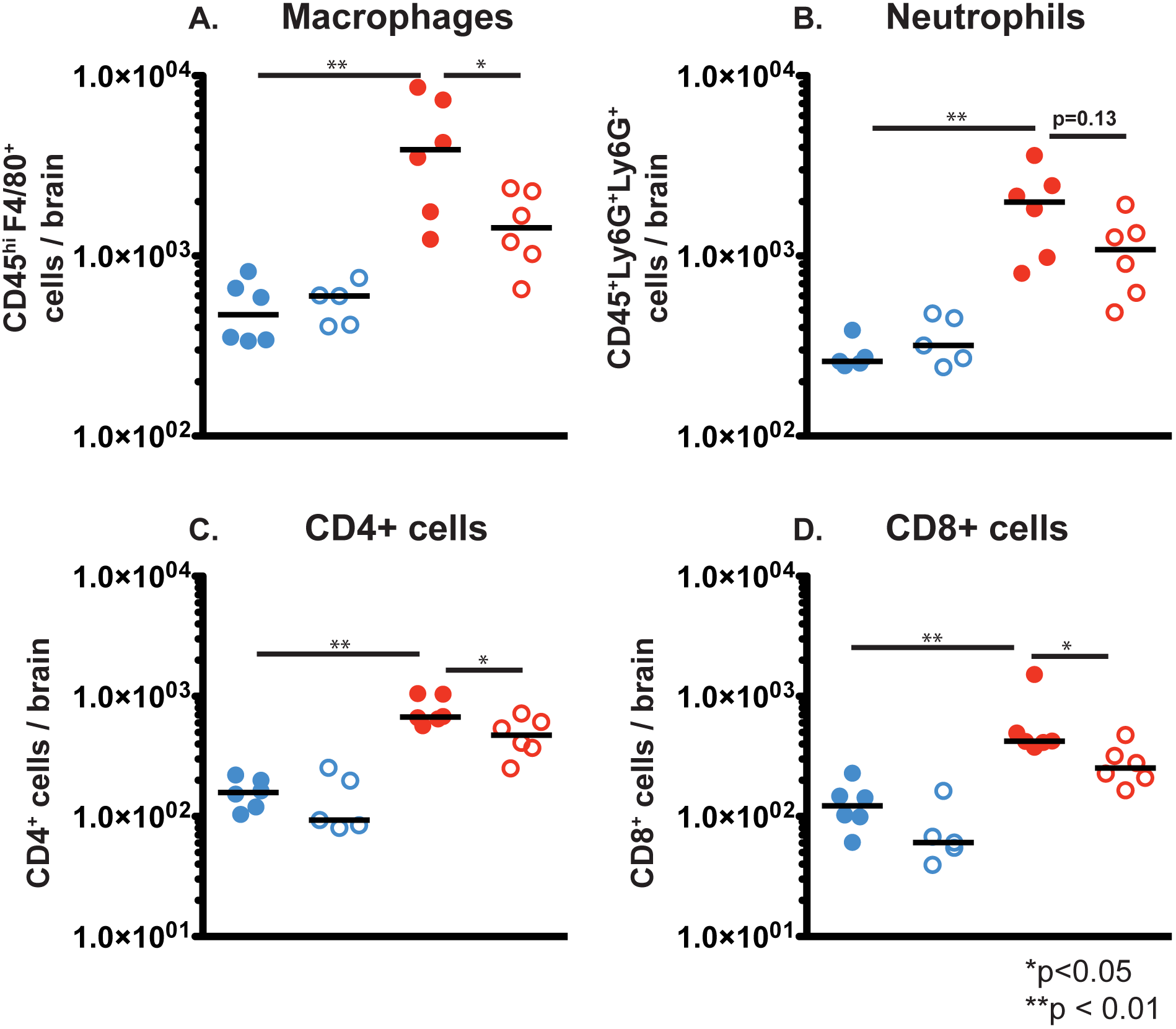
Administration of purified GXM to mice inoculated intracranially with acapsular *C. neoformans* reduces brain immune infiltration. These data are from the same experiments as **Fig. 10**. Brain infiltrating immune cells were detected by flow cytometry and broken into **(A)** CD45^hi^F4/80^+^ macrophages, **(B)** CD45^+^Ly6G^+^Ly6C^+^ Neutrophils, **(C)** CD4^+^ (T cells), **(D)** CD8^+^ (T cells). P-values were calculated using a Mann-Whitney test.

## Methods

### Conditioned media collection

*C. neoformans* cells were cultured overnight in YNB+2%glucose at 30 °C before subculturing 1:100 in the desired medium. Culture OD_600_ readings were taken 24 hours later and were normalized to the lowest measured OD_600_. Cells were pelleted by centrifugation at 3000xg for 5 min. The supernatant was collected and passed through a 0.22 μm filter, yielding conditioned media.

The following growth media were used in this study: YPAD (20g/L bacto-peptone, 10g/L bacto-yeast extract, 2% glucose, 0.4g/L adenine sulfate). YPD (20g/L bacto-peptone, 10g/L bacto-yeast extract, 2% glucose); YNB (Difco REF 291940) +2% glucose; 25% YNB+2% glucose; Low iron media (LIM) (5g/L asparagine, 0.4g/L K_2_HPO_4_, 0.1g MgSO_4_·7H_2_0, 0.5mg/L thiamine, 0.029mg/L boric acid, 1.88mg/L CuSO_4_·5H_2_0, 0.36mg/L MnCl_2_·4H_2_O, 0.021mg/L ZnCl_2_, 0.18mg/L NaMoO_4_·2H_2_0, 0.05mg/L CaCl_2_·2H_2_0, 0.05mM bathophenanthroline disulfonic acid (BPDS), 1mM EDTA, 2% glucose, 50mM MOPS pH 6.0), 10% Sabouraud’s dextrose (Difco REF 238230) buffered with 50mM HEPES pH 8.0, HEPES pH 7.3, MOPS pH 6.0, or MES pH 5.0; YCB (Difco REF 239110) +5g/L urea; YCB+0.5g/L urea.

### Conditioned media blots

10 μl of conditioned media collected from *C. neoformans* cultures were loaded into a 0.6% agarose gel and run at 33V for 18-20 hours at 0.5X TBE. The gels were processed with a 10 minute depurination rinse in a 0.25M HCl solution, followed by a 30 minute denaturation incubation in a 1.5M NaCl/0.5M NaOH solution, and a 30 minute neutralization incubation in 1.5M NaCl/0.5M Tris-HCl, pH 7.5. The gels were rinsed in distilled water following each incubation. Gel contents were subsequently transferred to a positively charged membrane using a standard Southern blot protocol with 10X SSC (saline-sodium citrate) in the reservoir. After overnight transfer, the blots were soaked briefly in 2X SSC and dried. Blots were then blocked for 1 hour in 1X PBS+5% milk and incubated shaking overnight at 4 °C in 1X PBS+5% milk with 1:40,000 anti-GXM monoclonal antibody. The following morning, blots were rinsed 3 times in 1X PBS, incubated 2 hours in 1X PBS+5% milk with 1:2500 goat anti-mouse HRP antibody, and washed for 2.5 hours in 1X PBS+0.1% tween-20, changing the wash buffer every 20 minutes. For imaging, blots were developed with Clarity Western ECL substrate (BioRad Cat. 170-5061) and imaging on a BioRad Western Blot Imager. Anti-GXM monoclonal antibodies used in this study: F12D2, 1326 (Thomas Kozel, University of Nevada, Reno).

### Cell measurements

*C. neoformans* cells collected from laboratory media were spun down at 3000xg for 5 min, washed twice in 1X PBS and resuspended in 1X PBS. To collect cells from infected mouse organs, 1 mL of organ homogenate was passed through a 70 μm cell strainer (Fisher Cat. No. 22-363-548). At this junction, capsule measurement methods were the same for both laboratory-grown and mouse-isolated *C. neoformans* cells. Cells were fixed for 15 minutes in 2% paraformaldehyde before washing twice with 1X PBS, and resuspending in 100 μl of 1X PBS. 4 μl of cell suspension was mixed with 4 μl of india ink (Higgins No. 44201) on a microscope slide, coverslipped and visualized. Successive, representative pictures were taken from the outside of the coverslipped area toward the middle, because smaller cells tended to spread towards the edges of the coverslip more so than larger cells. Total cell diameter was measured as the distance from one edge of the capsule to the opposite edge. Cell body diameter was measured as the distance from one edge of the cell wall to the opposite edge. Capsule thickness was calculated as the total cell diameter, minus the cell body diameter, and divided by two; (total cell diameter–cell body diameter)/2.

### Screen for exo-GXM mutants

Cells were spotted from 96 well frozen stocks to omnitrays containing YPD agar, then grown for 48 hours at 30°C. Colonies are used to inoculate deepwell plates containing 1 ml yeast nitrogen base (YNB) per well. Deepwell plates were grown at 37°C for 48 hours with shaking (280 rpm). 10 μl of YNB culture were then used to inoculate 10% Sabouraud’s (pH 7.3) cultures, which were then grown at 37°C for 48 hours with shaking. After growth, all cultures, either YNB or 10% Sabouraud’s, pH 7.3, were harvested by centrifugation, then the supernatant was collected and stored for analysis.

We analyzed exo-GXM in YNB supernatants by dot blotting 4 μl of supernatant into each well of a dot blotter containing positively charged nylon membrane pre-soaked in 2X SSC, then applying vacuum. Membranes were air dried, then blocked and incubated with anti-GXM F12D2 antibody using standard procedures (see Materials and Methods section: *Conditioned media blots*). 10% Sabouraud’s conditioned media samples were run on agarose gels and transferred to nylon membranes (see Materials and Methods section: *Conditioned media blots*).

Once we identified mutants with altered exo-GXM levels (decreased in YNB cultures or increased in 10% Sabouraud’s, pH 7.3 cultures, we grew all mutants in 10% Sabouraud’s, pH 7.3, then measured capsule thickness. Mutants with decreased cell surface capsule thickness (approximately 25% decrease compared to wild-type cells) were eliminated from further analysis. We then repeated the growth and exo-GXM blot for each strain. We normalized for cell density (to account for slow growing mutants), filtered the conditioned medium through a 0.22 μm filter to remove cells, and ran 10 μl of conditioned medium on an agarose gel using the procedure described in (see Materials and Methods section: *Conditioned media blots*). Finally, we stained for exposure of PAMPs such as chitin and mannoprotein (see Materials and Methods section: *Lectin Staining*) and removed mutants with increased exposure.

### Lectin Staining

Cells grown for 24 hours in the appropriate media were pelleted, washed twice in 1X PBS and fixed for 12 minutes in 2% paraformaldehyde. Cells were then washed twice in 1X PBS and resuspended in 1X PBS. To an aliquot of cells, wheat germ agglutinin (WGA) conjugated to fluorescein (Vector Labs Cat. No. FL-1021) was added to a final concentration of 5 μg/ml, and incubated 30 minutes at room temperature with shaking. At the end of the WGA incubation, concanavalin A (ConA) conjugated to rhodamine (Vector Labs Cat. No. RL-1002) was added to a final concentration of 50 μg/ml. Cells were wash once in 1X PBS and imaged immediately.

### Melanization and urease secretion

Cells grown overnight in YNB were washed twice in 1X PBS and resuspended to a final concentration of 2.5x10^6^ cells/mL in 1X PBS. 10μl of cell suspension was spotted onto L-DOPA containing agar or Christensen’s urea agar (Sigma 27048). Plates were checked daily for changes in melanization (brown/black colonies on L-DOPA), and urease secretion (pink coloration surrounding colonies on Christensen’s urea).

### GXM purification

GXM was purified as described previously (30). Briefly, 100 mL *C. neoformans* cells were cultured in YNB + 2%glucose for 5 days at 30°C. Cultures were centrifuged at 12,000xg for 15min and the supernatant collected. Polysaccharides were precipitated from the supernatant overnight with the addition of 3 volumes of 95% EtOH at 4 °C. The solution was then centrifuged at 15,000xg for one hour, resuspended in 0.2M NaCl and sonicated. After sonication, 3mg hexadecyltrimethylammonium bromide (CTAB) (Fisher Cat. No. 227160) per 1 mg precipitate was slowly added to the solution on low heat. After removing from heat, another 2.5 volumes of 0.5mg CTAB was added. The solution was centrifuged at 11,000xg for 2 hours, and the pellet washed in 10% EtOH to remove any remaining CTAB. After an additional centrifugation at 18,000xg, the pellet was resuspended in 1M NaCl and sonicated for 2 hours. Once the GXM was solubilized, it was dialyzed (3.5kDa cutoff) versus sterile distilled water and then lyophilized. Purified, lyophilized GXM was stored at -80°C for subsequent use.

### Adherence assay

We used a slightly modified protocol of biofilm formation and 2,3-Bis-(2-Methoxy-4-Nitro-5-Sulfophenyl)-2*H*-Tetrazolium-5-Carboxanilide (XTT) analysis, as described previously (46, 47). Briefly, 5 mL cultures were grown overnight in YNB+2%glucose at 30 °C, pelleted, washed in 1X PBS, and resuspended in 1X PBS.

Cells were counted on a hemocytometer, diluted to 10^7^ cells/mL in the appropriate media and plated in 100 μl volumes in 96 μl polystyrene plates (avoiding edge wells). Sterile media was plated as a negative control. Plates were incubated for 48 hours at 37 °C to allow for adherence and biofilm maturation. Plates were then washed 3 times with 200 μl of 1X PBS+0.05% tween-20 using a BioTek 405 TS microplate washer set to an intermediate flow rate. To determine the relative levels of cells that remained after washing, we used the XTT reduction assay to quantitate metabolic activity as a proxy for viable cell density. After plate washing, 100 μl of a solution containing 0.5g/L XTT (Fisher Cat. No. X6493) and 4 μM menadione (Sigma Cat. No. 58-27-5) in acetone in 1X PBS was added to each well. Menadione was added to fresh XTT solution immediately prior to adding the solution to a plate. Plates were incubated for 5 hours before moving 80 μl supernatant aliquots to a new plate to read absorbance at 490nm.

### Mice

For the intranasal infection model, we used ∼8-week-old female C57BL/6NJ mice (Jackson Labs). *C. neoformans* cells were harvested from overnight 30°C YPD cultures, washed two times in 1X PBS, resuspended in 1X PBS, and then counted with a hemocytometer to determine the inoculum. Mice were anesthetized with ketamine/dexdomitor (mg/g) intraperitoneally before suspending them on horizontally tied thread by their front incisors. Mice were kept warm with a heat lamp and inoculated intranasally with 2.5x10^4^ *C. neoformans* cells in 50 μl 1X PBS. After 10 minutes, mice were removed from thread and administered the reversal agent antisedan (∼0.0125mg/g) intraperitoneally. For survival analyses, mice were weighed daily and euthanized by CO_2_ asphyxiation and cervical dislocation, when they lost 15% of their initial mass. Mice used to analyze fungal burden, capsule size, and GXM levels were euthanized by the same measures at designated time points. Mice used for flow cytometry analysis were anesthetized with isoflurane and intracardially perfused with cold 1X PBS before cervical dislocation and brain extraction.

For the intracranial infection model, we used ∼6-week-old female C57BL/6NJ mice (Jackson labs). *C. neoformans* inoculum was prepared as described above. Prior to inoculation, mice were anesthetized with ketamine/dexdomitor, as above. Mice were inoculated intracranially with 200 *C. neoformans* cells in 30μl 1X PBS via a 26Gx1/2 needle. Animals were then administered antisedan to speed recovery.

### Fungal Burden

Organs were harvested from euthanized mice, placed on ice, and homogenized with a Tissue Master Homogenizer (Omni International) in 5 mL 1X PBS. Serial dilutions of organ homogenates were plated on Sabouraud’s dextrose agar with 10mg/mL gentamycin and 100 mg/mL carbenicillin, and stored at 30°C in the dark for three days. Resulting colony forming units (CFU) were then counted to determine fungal burden.

### GXM ELISA

500 μl of the same mouse organ homogenate used for CFU counts and *C. neoformans* cell measurements was collected and spun down at 3,000g for 5 minutes. The supernatant was then passed through a 0.22 μm filter to remove cells. GXM levels in the resulting were quantified using the ALPHA Cryptococcal Antigen enzyme immunoassay (IMMY Ref. CRY101). GXM purified from *C. neoformans* cultures was diluted to generate standard curves.

### Histology

Perfused mouse brains were divided in half and fixed overnight in 4% paraformaldehyde. 8 μm thick sagittal slices were mounted on microscope slides and stored at -20 °C. Successive sections were stained with hematoxylin and eosin or Grocott’s methenamine silver (ThermoFisher Scientific Cat. No. 87008).

### Flow cytometry

Perfused mouse brains were collected in RPMI, ground gently to disperse tissue and spun in a 90% Percoll (Sigma Cat. No. P1644) with a 63% Percoll underlay to isolate leukocytes at the interface. Leukocytes were resuspended in FACS buffer (1X PBS, 1% bovine serum albumin), and stained with the appropriate fluorescently labeled antibodies. Labeled cells were fixed for 20 minutes in 4% paraformaldehyde before analysis on a LSRFortessa (BD Biosciences). Antibodies used in this study (eBiosciences): CD45-efluor450 (48-0451-82), CD4-APC (Cat. No. 17-0041-82), CD8-FITC (11-0081-82), F4/80-FITC (11-4801-82), Ly6G-FITC (11-5931-82), Ly6C-APC (17-5932-82).

